# Impaired state-dependent potentiation of GABAergic synaptic currents triggers seizures in an idiopathic generalized epilepsy model

**DOI:** 10.1101/2020.05.10.087114

**Authors:** Chun-Qing Zhang, Mackenzie A. Catron, Li Ding, Caitlyn M. Hanna, Martin J. Gallagher, Robert L. Macdonald, Chengwen Zhou

## Abstract

Idiopathic generalized epilepsy(IGE) patients have genetic causes and their seizure onset mechanisms particularly during sleep remain elusive. Here we proposed that sleep-like slow-wave oscillations(0.5 Hz SWOs) potentiated excitatory or inhibitory synaptic currents in layer V cortical pyramidal neurons from wild-type(wt) mouse *ex vivo* brain slices. In contrast, SWOs potentiated excitatory, not inhibitory, currents in cortical neurons from heterozygous(het) knock-in(KI) IGE mice(GABA_A_ receptor γ2 subunit *Gabrg2^Q390X^* mutation), creating an imbalance between evoked excitatory and inhibitory currents to effectively prompt neuronal action potentials. Similarly, more physiologically similar up/down-state(present during slow-wave sleep) induction in cortical neurons could potentiate excitatory synaptic currents within slices from wt/het *Gabrg2^Q390X^* KI mice. Consequently, SWOs or up/down-state induction *in vivo* (using optogenetic method) could trigger epileptic spike-wave discharges(SWDs) in het *Gabrg2^Q390X^* KI mice. To our knowledge, this is the first operative mechanism to explain why epileptic SWDs preferentially happen during non-REM sleep or quiet-wakefulness in human IGE patients.

## Introduction

Epilepsy affects more than 3 million people in the United States and causes cognitive and other comorbidities. In two thirds of patients the causes are unknown (possible genetic origins, termed idiopathic generalized epilepsy[IGE]). The Epi4K consortium has identified many new genetic mutations in IGE patients(Allen et al., 2013; Striano & Zara, 2017), making it imperative for researchers to elucidate seizure onset mechanisms in patients with different genetic mutations and varied epileptic symptoms. While it’s widely acknowledged that epileptic events randomly occur in patients, including IGE patients, epileptic spike-wave discharges(SWDs) do preferentially appear during slow-wave sleep or quiet-wakeful states(motor immobility)(Ahmed & Vijayan, 2014; F. Arain, Zhou, Ding, Zaidi, & Gallagher, 2015; Bagshaw, Rollings, Khalsa, & Cavanna, 2014; Halasz, Terzano, & Parrino, 2002; Ng & Pavlova, 2013; Shouse, Scordato, & Farber, 2004). During slow-wave sleep or quiet-wakeful states, brain activity in EEG recordings exhibits large amplitude alteration at low delta frequency (around 0.5 Hz, slow-wave oscillations[SWOs])(Demanuele, Sonuga-Barke, & James, 2010; Lagarde et al., 2019; Petersen, Hahn, Mehta, Grinvald, & Sakmann, 2003) and cortical neurons can undergo homeostatic changes of synaptic currents and ion channels(Kurotani, Yamada, Yoshimura, Crair, & Komatsu, 2008; Liu, Faraguna, Cirelli, Tononi, & Gao, 2010; Turrigiano, 2008). Thus this will lead to the dynamic interaction between synaptic currents in cortical neurons(Dehghani et al., 2016; Haider, Duque, Hasenstaub, & McCormick, 2006; Shu, Hasenstaub, & McCormick, 2003), which might play a role in seizure onset mechanisms in IGE patients.

Our groups have been working on GABA_A_ receptor mutations Q390X in γ2 (*Gabrg2^+/Q390X^*) and α1(*Gabra1^+/A322D^*) subunits identified in human IGE patients including Dravet syndrome(Kang, Shen, Zhou, Xu, & Macdonald, 2015; Warner, Smith, & Kang, 2019). Human patients exhibit epileptic SWDs during non-rapid eye-movement sleep (NREM sleep)(Bagshaw et al., 2014; Halasz, Kelemen, & Szucs, 2013; Verbeek et al., 2015) and some IGE patients are resistant to conventional antiepileptic drug treatments. These mutations expressed in heterozygous(het) knock-in(KI) mice(heterozygosity simulates human patient conditions) can cause mice to generate febrile seizures, generalized epilepsy(*Gabrg2^+/Q390X^*)(Kang & Macdonald, 2016), absence epilepsy (both *Gabrg2^+/Q390X^* and *Gabra1^+/A322D^*)(F. M. Arain, Boyd, & Gallagher, 2012; Ding et al., 2010) and juvenile myoclonic seizures (*Gabra1^+/A322D^*)(F. Arain et al., 2015; Ding et al., 2010; Warner et al., 2019). Moreover, epileptic SWDs in these het KI mice are present while no ostensible motor behaviors are observed(F. M. Arain et al., 2012; Ding et al., 2010; Kang & Macdonald, 2016), similar to quiet-wakeful states. In addition, both mutations cause dysfunctional GABA_A_ receptor subunit aggregation inside cells(Kang, Shen, Lee, Gallagher, & Macdonald, 2010; Zhou et al., 2013), which would influence synaptic plasticity. Thus, we hypothesized that SWOs during NREM sleep and quiet-wakeful state could generate state-dependent dysfunctional plasticity of synaptic currents, particularly inhibitory currents, and eventually trigger seizure onset in these mouse IGE models.

In this study we found that sleep-like SWOs(0.5 Hz) or up/down-state could induce synaptic plasticity in cortical pyramidal neurons(layer V), and that state-dependent potentiation of excitatory or inhibitory synaptic currents remained intact in cortical neurons from wild-type(wt) littermates. In contrast, only excitatory synaptic currents, but not inhibitory, could be potentiated in neurons from het *Gabrg2^+/Q390X^* KI mice, creating unbalanced excitation and inhibition to effectively elicit action potentials in neurons. With *in vivo* up/down-state induction by optogenetic manipulation(simulating *ex vivo* SWOs), epileptic SWDs could be triggered with significantly higher incidence and longer duration being accompanied by animal immobility and other epileptic behaviors in het *Gabrg2^+/Q390X^* KI mice, suggesting that SWOs or up/down-states *in vivo* can trigger epileptic seizure onset in IGE patients while they are in NREM sleep or quiet-wakefulness state.

## Results

To address the roles of SWOs on epileptic SWD generation, we used wt littermate and het *Gabrg2^+/Q390X^* KI mice, and transgenic wt or het *Gabrg2^+/Q390X^* KI mice expressing Thy1-halorhodopsin protein in cortical neurons for optogenetic SWO induction *ex vivo* or *in vivo*. The expression of Thy1-halorhodopsin in neurons was confirmed by enhanced green fluorescent protein(EGFP) expression in cortical neurons and physiology experiments *ex vivo*. Moreover, using a method by Kurotani et al.(Kurotani et al., 2008), layer V cortical pyramidal neurons within somatosensory cortex(controlling cortical epileptic activity propagation(Polack et al., 2007; Wester & Contreras, 2012)) were injected with sinusoidal currents(at 0.5 Hz, current-clamp) to cause neuronal membrane potential oscillating between resting membrane potential −70 - −75 mV and −50 mV, very similar to neuron up/down-state alteration during NREM sleep(Petersen et al., 2003; Steriade, Timofeev, & Grenier, 2001), which was defined as SWOs in this study. In addition, cortical neurons from wt littermates (n=33 for each, capacitance 108.95 ± 6.67 pF; input resistance 87.87 ± 7.75 MΩ, resting membrane potential −72.07 ± 0.57 mV, action potential amplitude 79.64 ± 1.54 mV[measured from precedent resting membrane potential]) were not significantly different from those neurons from het *Gabrg2^+/Q390X^* KI mice in basic physiology properties(n=37 for each, capacitance 106.89 ± 4.72 pF, t-test p=0.798; input resistance 77.47 ± 6.63 MΩ, t-test p=0.309; resting membrane potential −71.52 ± 0.54 mV, t-test p=0.491; action potential amplitude 82.84 ± 1.35 mV, t-test p=0.119).

### SWO induction state-dependently potentiates excitatory synaptic currents in layer V cortical neurons from wt and het *Gabrg2^+/Q390X^* KI mice

α-amino-3-hydroxy-5-methyl-4-isoxazolepropionic acid receptor(AMPAR)-mediated spontaneous excitatory synaptic currents (sEPSCs) were isolated by clamping cortical neurons at chloride anion reversal potentials (−55.8 mV) to examine whether SWO induction could potentiate sEPSCs in cortical neurons from brain slices *ex vivo*. Compared with pre-SWO baseline sEPSCs, SWO induction (0.5 Hz) for 10 min significantly increased sEPSC amplitudes with stable access resistances(Fig. 1A-C) in neurons from wt littermates. Potentiated sEPSCs reached their peak around 10 min after SWO induction and then decayed to a plateau around 30 min, remaining larger than baseline sEPSC amplitudes(Fig. 1B). Both summary data of pre- and post-SWO sEPSC amplitudes(Fig. 1C inset, from 16.69 ± 1.85 to 27.90 ± 4.13 pA, n=8 cells, n=5 mice, paired t-test p=0.004) and cumulative sEPSC amplitude distribution indicated that SWO induction did significantly increase AMPAR-mediated sEPSCs in neurons from wt littermates(Fig. 1C, K-S test p=0.00001). Similar to sEPSC potentiation in neurons from wt littermates, post-SWO sEPSCs in cortical neurons from het *Gabrg2^+/Q390X^* KI mice also exhibited significant enhancement (from 18.89 ± 1.52 to 28.12 ± 3.47 pA, n=9 cells, n=6 mice, paired t-test p=0.009) during post-SWO period (Fig. 1D-F, K-S test p=0.00001), indicating that SWO induction could also potentiate sEPSCs in cortical neurons from het *Gabrg2^+/Q390X^* KI mice. Moreover, we did not observe any significant changes between pre- and post-SWO sEPSC frequency and sEPSC decay in neurons from wt littermates (wt [n=8 cells, n = 5 mice], frequency from 2.43 ± 0.69 to 1.71 ± 0.36 Hz, paired t-test p = 0.36; decay τ, from 18.57 ± 5.35 to 25.13 ± 3.89 ms, paired t-test p = 0.304) or het *Gabrg2^+/Q390X^* KI mice([n=9 cells, n=6 mice], frequency from 1.67 ± 0.25 to 3.06 ± 0.84 Hz, paired t-test p=0.10; decay τ from 12.80 ± 1.62 to 34.15 ± 10.89 ms, paired t-test p = 0.078).

**Figure 1.**
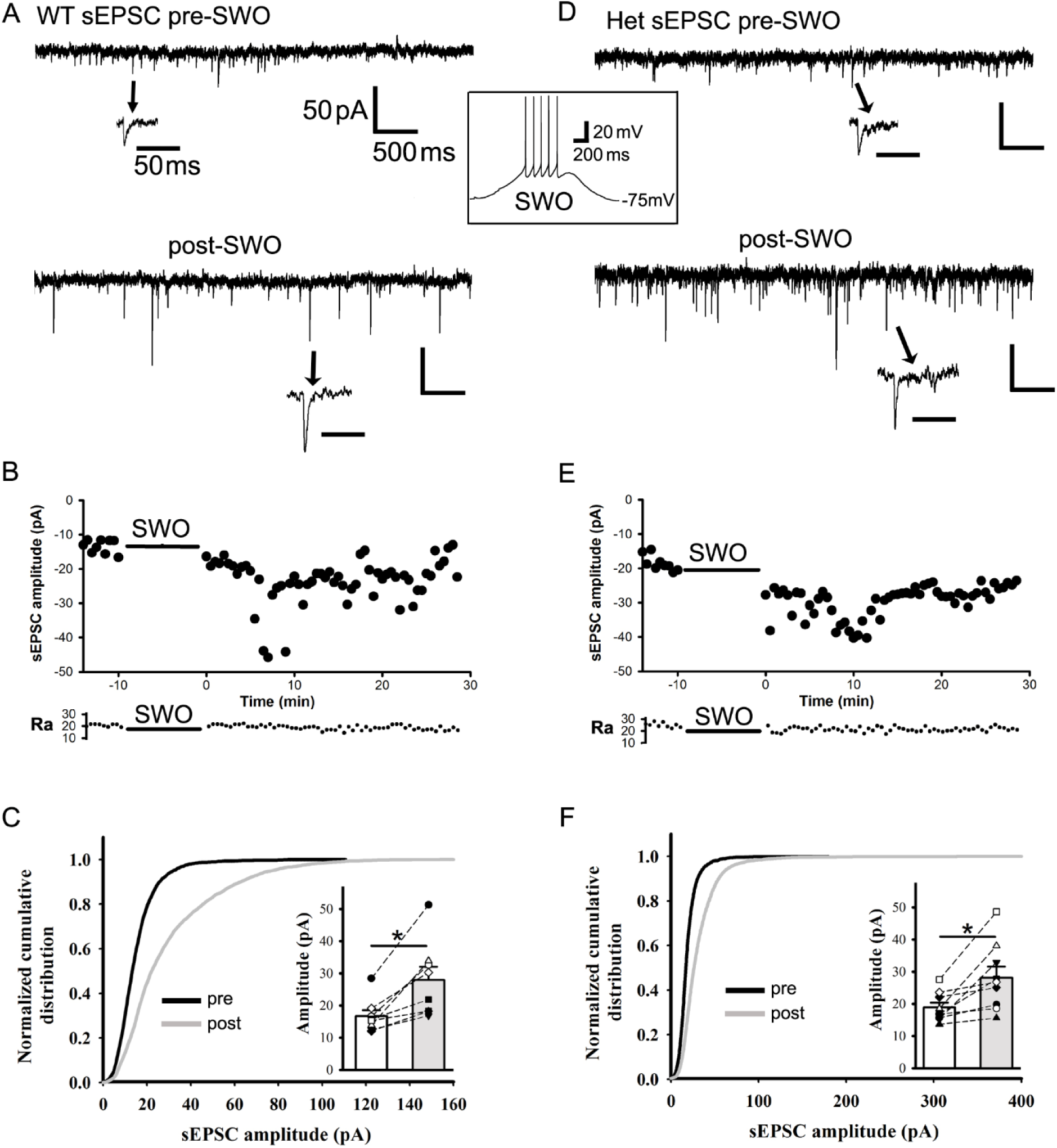
SWO induction potentiates spontaneous excitatory synaptic currents in layer V cortical pyramidal neurons from wt or het *Gabrg2^+/Q390X^* KI mice. Panels A and D are representative traces pre-SWO (top) and post-SWO sEPSCs (lower) for wt littermates (left column) and het *Gabrg2^+/Q390X^* KI mice(right column). The arrows point to individual sEPSC events in extended time scales. The middle inset is one representative SWO induction trace from rest membrane potentials (−75 mV). Scale bars are indicated as labeled. Panels B and E show time courses of pre-SWO and post-SWO sEPSC amplitudes for corresponding recordings from panel A(wt) and D(het). Each data point in the panels B/E was obtained by averaging all sEPSC events during continuous 5 s recordings. The lower panel of access resistance (Ra) shows its accompanied values during the whole 42.5 min recordings. Panels C and F show normalized cumulative histogram of sEPSC events for wt and het *Gabrg2^+/Q390X^* KI mice. Insets are summary data of pre-SWO and post-SWO sEPSC values (n=8 neurons, n=5 mice, paired t-test p= 0.004 for wt; n=9 neurons, n=6 mice for het *Gabrg2^+/Q390X^* KI mice, paired t-test p=0.009)(wt cumulative curves used 4905 synaptic events for pre-SWO[during 4.2 min baseline recordings for each neuron, total n=8 neurons] and 4506 synaptic events for post-SWO [during final 4.2 min recordings for each neurons, total n=8 neurons], while het cumulative curves used 3886 synaptic events for pre-SWO[during 4.2 min baseline recordings for each neuron, total n=9 neurons] and 9519 synaptic events for post-SWO[during final 4.2 min recordings for each neuron, total n=9 neurons]. Each data point for inset graphs was obtained by averaging sEPSC events during baseline or post-SWO last 4.2 min recordings).

In addition to the synaptic scaling up by low-level neuron activity (0.5 Hz), we also tested synaptic scaling down by high-level neuron activity(50 Hz). When neuron activity was elevated with 50 Hz oscillating currents injected, cortical neurons from either wt littermates or het *Gabrg2^+/Q390X^* KI mice exhibited dramatic attenuation in sEPSC amplitudes (supplemental Fig. S1) for more than 30 min compared to baseline sEPSCs(wt and het data combined[n=6 cells, n=4 mice], from 14.22 ± 0.77 to 10.00 ± 0.76 pA, paired t-test p=0.004) and showed no changes in sEPSC frequency (2.34 ± 0.49 Hz to 1.73 ± 0.14 Hz, paired t-test p=0.218) and in sEPSC decay(decay τ, from 15.40 ± 4.48 to 21.88 ± 4.62 ms, paired t-test p = 0.285). This suggested the possibility of homeostatic plasticity mechanism in our study(Turrigiano, 2008; Turrigiano & Nelson, 2004), although we did not use same TTX/NBQX or picrotoxin to manipulate neuron activity in culture neurons as other studies(Turrigiano & Nelson, 2004). Thus, we further assessed whether this state-dependent potentiation of sEPSCs(scaling up by 0.5 Hz) involved a homeostatic plasticity mechanism by using one retinoid acid synthesis blocker 4-(diethylamino)-benzaldehyde(DEAB)(Aoto, Nam, Poon, Ting, & Chen, 2008; Chen, Lau, & Sarti, 2014; Li, Park, Zhong, & Chen, 2019; Turrigiano & Nelson, 2004), low-level BAPTA-AM(15 µM, maintaining low-level Ca^2+^)(Ibata, Sun, & Turrigiano, 2008; Wang, Zhang, Hintze, & Chen, 2011), KN93(βCaMKII antagonist)(Thiagarajan, Piedras-Renteria, & Tsien, 2002) and nifedipine(L-type calcium channel blocker)(Kurotani et al., 2008)(supplemental Fig. S2). These chemical agents can either block(DEAB and KN93) or enhance(low-level Ca^2+^ and nifedipine) homeostatic synaptic plasticity in cortical neurons. With 40 µM DEAB in ACSF, SWO induction could not enhance any sEPSCs in cortical neurons from either wt littermate or het *Gabrg2^+/Q390X^* KI mice(supplemental Fig. S2A and 2B[n=10 cells, n=4 mice for each comparison], amplitude from 15.05 ± 1.48 to 15.52 ± 2.30 pA, paired t-test p=0.632; frequency from 4.45 ± 0.96 to 4.17 ± 1.00 Hz, paired t-test p=0.733; decay τ, from 15.50 ± 1.96 to 27.80 ± 8.15 ms, paired t-test p = 0.112), confirming that homeostatic synaptic plasticity did mediate SWOs-induced potentiation of sESPCs in cortical neurons from wt and het *Gabrg2^+/Q390X^* KI mice. Similar to DEAB suppressing action, administration of KN-93(2 µM) inside cells also suppressed the sEPSC potentiation in cortical neurons from either wt or het mice (supplemental Fig. S2B[n=7 cells, n=3 mice], sEPSC amplitude from 12.78 ± 1.17 to 11.04 ± 1.04 pA, paired t-test p=0.013; no change in sEPSC frequency from 2.91 ± 0.92 to 1.41 ± 0.36 Hz, paired t-test p=0.117, while longer decay τ 9.75 ± 1.45 to 25.02 ± 3.88 ms, paired t-test p=0.005). However, treatment brain slices with BAPTAM-AM(15 µM) (at least 40 min) or addition of nifedipine (20 µM) in ACSF could enhance potentiation of sEPSCs in cortical neurons from either wt or het *Gabrg2^+/Q390X^* KI mice(BAPTA-AM [n=5 cells, n=3 mice], sEPSC amplitude from 9.55 ± 0.56 to 12.99 ± 0.88 pA, paired t-test p=0.006; sEPSC frequency from 0.86 ± 0.20 to 1.88 ± 0.41 Hz, paired t-test p=0.021, while longer decay τ 10.98 ± 2.28 to 26.80 ± 2.26 ms, paired t-test p=0.009)(nifedipine [n=7 cells, n=4 mice], sEPSC amplitude from 12.14 ± 1.21 to 21.49 ± 3.38 pA, paired t-test p=0.044; sEPSC frequency from 4.40 ± 0.93 to 3.03 ± 0.55 Hz, paired t-test p=0.063 and decay τ 15.99 ± 3.02 to 24.89 ± 3.61 ms, paired t-test p=0.105). All together, these pharmacological results did indicate the homeostatic plasticity involvement in sEPSC enhancement by 0.5 Hz SWOs in cortical neurons. In addition, treatment brain slices with only BAPTA-AM(15 µM) seemed to slight decrease baseline sEPSC amplitudes of cortical neurons, which may be related to action of BAPTA buffering intracellular calcium on transmitter release(Rozov, Burnashev, Sakmann, & Neher, 2001).

### SWO induction state-dependently potentiates inhibitory synaptic currents in cortical neurons from wt, but not from het *Gabrg2^+Q/390X^* KI mice

In cortical neurons, synaptic excitation and inhibition are always proportional/balanced to avoid cortical hyperexcitability(Shu et al., 2003; Haider et al., 2006; Dehghani et al., 2016). Thus, we reasoned whether this SWO induction could also potentiate GABAergic currents in cortical neurons. GABA_A_ receptor-mediated spontaneous inhibitory synaptic currents(sIPSCs) were isolated by using the AMPAR/kainate receptor antagonist NBQX (20 µM) in ACSF and neurons were voltage-clamped at −60 mV (chloride reversal potential −15 mV). SWO induction in cortical neurons significantly increased sIPSC amplitude compared to pre-SWO baseline from wt littermates (Fig. 2 A-C)(from 27.14 ± 3.79 to 39.06 ± 6.06 pA, n=10 cells, n=8 mice, paired t-test p=0.001; K-S test p=0.00001), similar to the potentiation of miniature IPSCs in neurons from Kurotani et al.(Kurotani et al., 2008). Moreover, the time course of potentiated sIPSCs exhibited a peak around 10 min after SWO induction(Fig. 2B), very similar to the time course of post-SWO sEPSCs in neurons from wt littermates(Fig. 1B). However, in neurons from het *Gabrg2^+/Q390X^* KI mice, SWO induction did not potentiate any sIPSCs compared to pre-SWO baseline (with stable access resistances) (Fig. 2D and E). Both summary data of pre- and post-SWO sIPSCs (Fig. 2 F inset, from 18.60 ± 2.08 to 16.86 ± 1.85 pA, n=11 cells, n=7 mice, paired t-test p=0.1076) and cumulative distribution analysis (Fig. 2F, K-S test p=0.061) confirmed no sIPSC potentiation in neurons from het KI mice, indicating that SWO-induced potentiation of sIPSCs in het KI mice was impaired. In addition, we did observe significant increase in post-SWO sIPSC frequency in neurons from wt littermates (from 3.41 ± 0.87 to 7.90 ± 1.70 Hz, n=10 cells, n=8 mice, paired t-test p=0.021), not from het KI mice (from 4.28 ± 1.20 to 2.45 ± 0.47 Hz, n=11 cells, n=7 mice, paired t-test p=0.148). Meanwhile, sIPSC decay in neurons exhibited longer decay after SWO induction from both wt and het *Gabrg2^+/Q390X^* mice (decay τ, wt from 10.18 ± 2.24 to 29.62 ± 7.86 ms, n=10 cells, n = 8 mice, paired t-test p = 0.0229; het from 10.46 ± 2.58 to 31.37 ± 8.09 ms, n=11 cells, n = 7 mice, paired t-test p = 0.0278), suggesting that GABAergic receptors might change their components after SWO induction.

**Figure 2.**
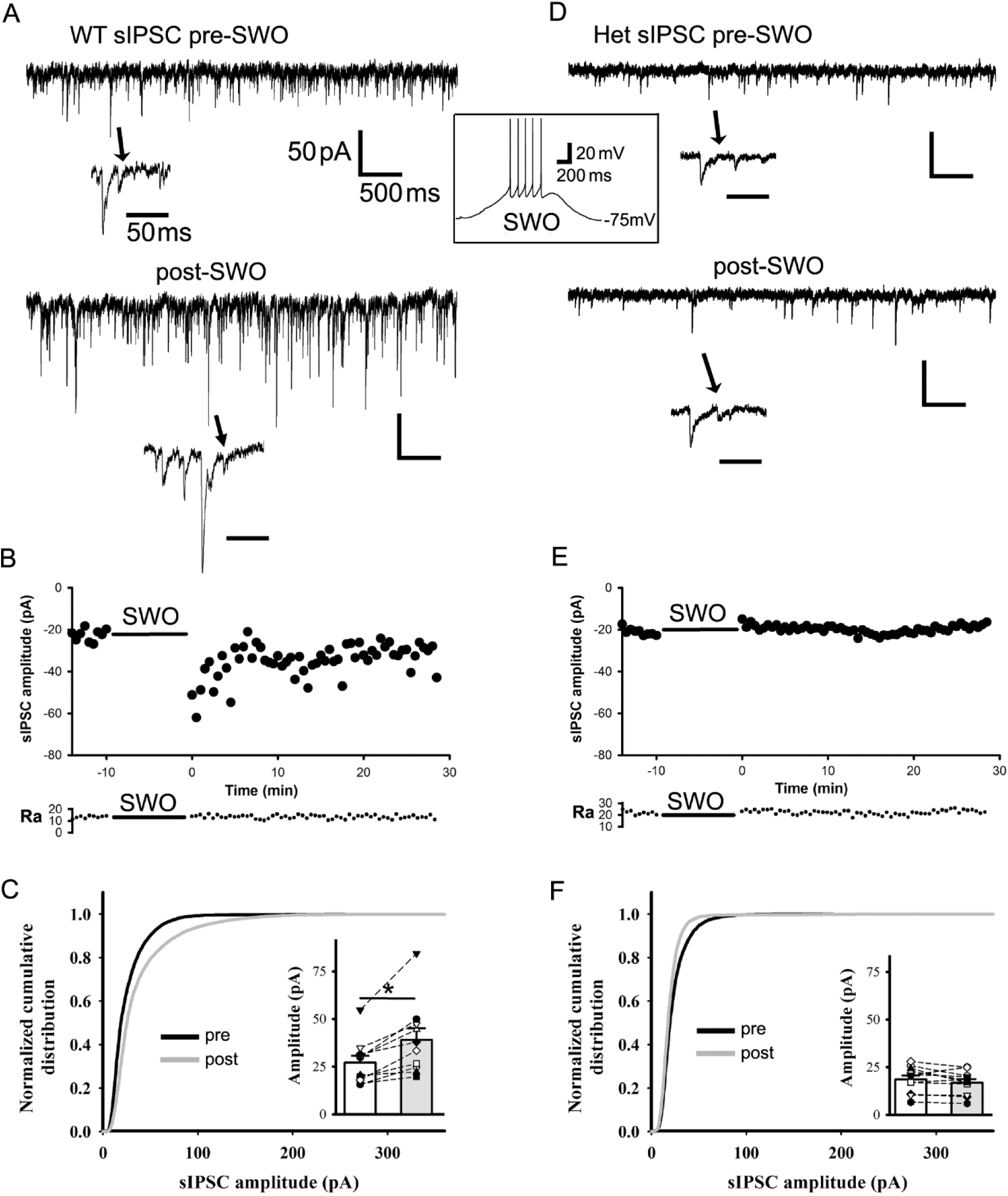
SWO induction potentiates spontaneous inhibitory synaptic currents in layer V cortical pyramidal neurons from wt littermates, but not from het *Gabrg2^+/Q390X^* KI mice. Panels A and D are representative traces pre-SWO (top) and post-SWO sIPSCs (lower) for wt littermates (left column) and het *Gabrg2^+/Q390X^* KI mice(right column). The arrows indicate individual sIPSC events in extended time scales. The middle inset is one representative SWO induction trace from resting membrane potentials (−75mV). Scale bars are indicated as labeled. Panels B and E show time courses of pre-SWO and post-SWO sIPSC amplitudes for corresponding recordings from panel A(wt) and D(het). Each data point in the panels B/E was obtained by averaging all sIPSC events during continuous 5s recordings. The low panel of access resistance (Ra) shows its values during the whole 42.5 min recordings. Panels C and F show normalized cumulative histogram of sIPSC events for wt and het *Gabrg2^+/Q390X^* KI mice. Insets are summary data of pre-SWO and post-SWO sIPSC values (n=10 neurons, n=8 mice, paired t-test p= 0.001 for wt; n=11 neurons, n=7 mice for het *Gabrg2^+/Q390X^* KI mice, paired t-test p=0.1076) (wt cumulative curves used 6293 synaptic events for pre-SWO[during 4.2 min baseline recordings for each neurons, total n=10 neurons] and 14299 events for post-SWO [during final 4.2 min recordings for each neuron, total n=10 neurons], while het cumulative curves used 7124 synaptic events for pre-SWO[during 4.2 min baseline recordings for each neurons, total n=11 neurons] and 6290 synaptic events for post-SWO[during final 4.2 min recordings for each neurons, total n=11 neurons]. Each data point for inset graphs was obtained by averaging sIPSC events during baseline or post-SWO last 4.2 min recordings).

We next examined the effect of elevated neuron activity on sIPSCs from wt littermates to determine whether this potentiation of sIPSCs in neurons involved homeostatic plasticity. After inducing 5 Hz membrane potential oscillation for 10 min, sIPSCs did undergo attenuation in amplitudes compared to baseline (supplemental Fig. S3 [n=5 cells, n=4 mice], amplitude from 20.37 ± 1.58 to 13.73 ± 1.11 pA, paired t-test p=0.004; frequency from 4.27 ± 1.14 to 1.60 ± 0.41 Hz, paired t-test p=0.048; sEPSC decay τ from 9.68 ± 1.17 to 23.36 ± 3.92 ms, paired t-test p = 0.041), suggesting that homeostatic plasticity of synaptic sIPSCs might involve SWO-induced sIPSC potentiation in neurons from wt littermates.

### Physiology-like up/down-state induction can potentiate excitatory synaptic currents in cortical neurons

Cortical neurons *in vivo* exhibit up/down-state alteration during slow-wave EEG oscillation during NREM sleep(Petersen et al., 2003; Steriade et al., 2001), essentially the same as SWO induction in brain slices *ex vivo*. As a surrogate for physiological up/down-state *in vivo*, we examined whether up/down-state induction within brain slices *ex vivo* could potentiate synaptic currents. Using a modified ACSF (containing 3.5 or 5 KCl, 1 Ca^2+^, 1 Mg^2+^ and 3.5 µM carbachol) and fast flow perfusion (6-7 mL/min)(Hajos & Mody, 2009; Neske, Patrick, & Connors, 2015; Sanchez-Vives & McCormick, 2000), we could induce long episodes(10s) of membrane depolarization(up-state, around −50 mV, with simultaneous action potential firing) from resting membrane potentials(down-state, around −70 mV) (Fig. 3A middle, n=7 cells, n=3 for wt and n=4 het *Gabrg2^+/Q390X^* KI mice). Compared to pre up/down-state baseline, up/down-state induction (10 min) significantly increased sEPSC in amplitudes(Fig. 3A-C[n=7 cells, n=7 mice], from 18.10 ± 2.19 to 27.69 ± 2.98 pA, paired t-test p = 0.003) and frequency (from 1.21 ± 0.22 to 2.52 ± 0.58 Hz, paired t-test p = 0.014) and slight longer decay τ(from 16.12 ± 1.68 to 51.01 ± 14.61 ms, paired t-test p = 0.069), which was very similar to results with SWO induction *ex vivo*. It was also noted that this modified ACSF within slices could induce many cortical neurons’ up/down-states at same time and changed the whole network activity to likely increase synaptic frequency. Furthermore, up/down-induced sEPSC potentiation within slices *ex vivo* could be blocked by DEAB treatment (40 µM DEAB, amplitude from 18.06 ± 1.96 to 16.14 ± 1.83 pA, n=6 cells, n=6 mice, paired t-test p=0.003) and no changes in frequency and decay τ ([n=6 cells, n=6 mice], frequency from 2.49 ± 0.78 to 2.18 ± 0.50 Hz, paired t-test p=0.691; decay τ from 21.73 ± 10.69 to 52.41 ± 11.63 ms, paired t-test p = 0.191). Together, these results suggested that up/down-state induction *in vivo* might use homeostatic plasticity mechanism.

**Figure 3.**
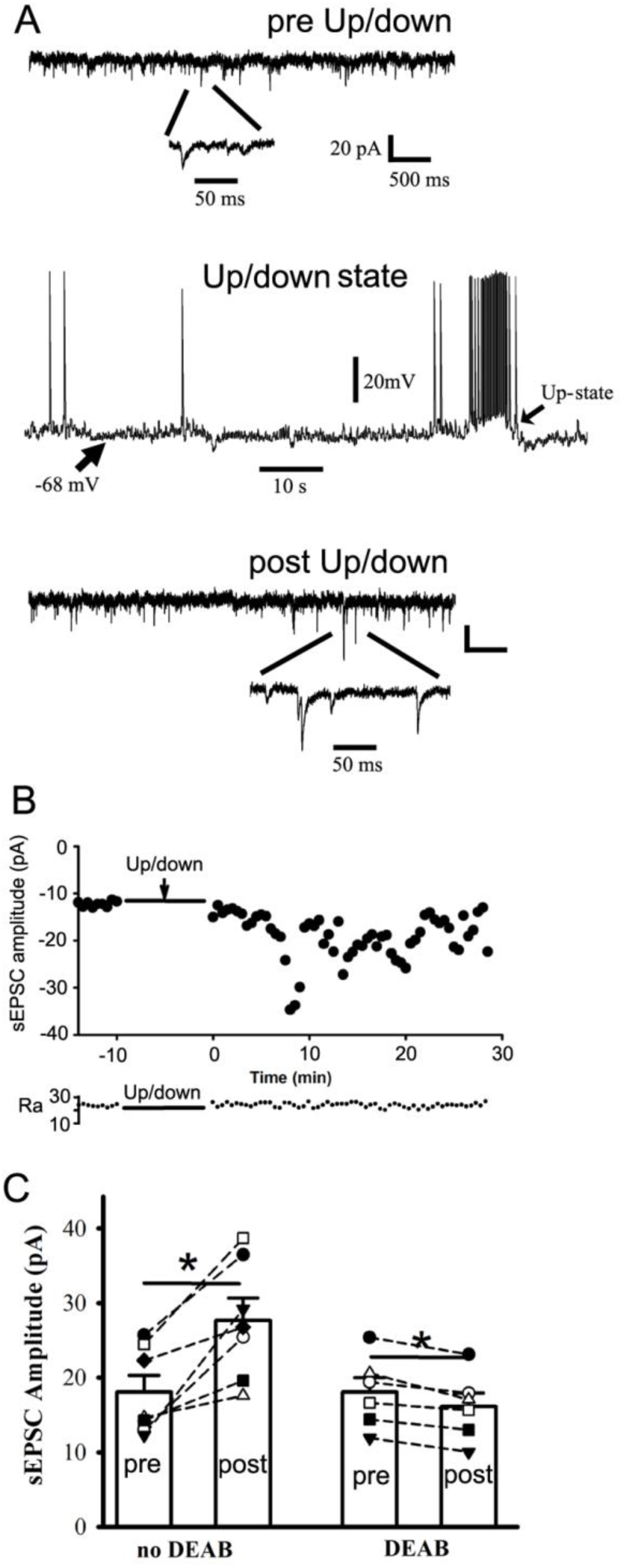
Physiologically similar up/down-state induction can potentiate excitatory synaptic currents in cortical neurons from wt or het *Gabrg2^+/Q390X^* KI mice. Panel A(top and bottom) shows representative traces for pre/post-up/down sEPSCs in cortical neurons. Below are expanded sEPSC events. Panel A(middle) shows one representative up/down-state induced by using fast flow perfusion (6-7ml/min) with a modified ACSF (containing [mM] 3.5 or 5 KCl, 1 Ca^2+^, 1 Mg^2+^ and 3.5 µM carbachol). Scale bars are indicated as labeled. Panel B shows time courses of pre/post-up/down-state sEPSC amplitudes for corresponding recordings from panel A. Each data point in these panels B was obtained by averaging all sIPSC events during continuous 5 s recordings. The lower part of panel B shows access resistance (Ra) values during the whole 42.5 min recordings. Panel C shows summary data of pre/post-up/down-state sEPSCs (n=7 neurons, n=7 mice, paired t-test p=0.003) and DEAB blockade effect (n=6 neurons, n=6 mice, paired t-test p=0.003)(each data point for panel C was obtained by averaging sEPSC events during baseline or post-up/down-state last 4.2 min recordings).

### Physiology-like up/down-state induction creates an imbalance between evoked inhibitory and excitatory synaptic currents in cortical neurons from het *Gabrg2^+/Q390X^* KI mice

It’s generally assumed that IPSCs and EPSCs in neurons from wt littermates keep in balance(Dehghani et al., 2016; Haider et al., 2006; Shu et al., 2003), suggesting that synaptic inhibition either equals or is larger than synaptic excitation. To assess the evoked(e) IPSC/EPSC balance in neurons from wt littermates and het *Gabrg2^+/Q390X^* KI mice, neurons were at clamped at −40 mV to simultaneously evoke both EPSCs and IPSCs and we compared their corresponding charge transfer as Nanou et al’s study[(Nanou, Lee, & Catterall, 2018), Fig 7 in their paper]. Compared with charge transfer ratio of wt control(1.38 ± 0.24, n=11, n=3 mice), wt up/down-state induction significantly increased the ratio of evoked(e)IPSC/eEPSC in cortical neurons(Fig. 4A/B, 3.64 ± 0.88, n=7 cells, n=3 mice, t-test p=0.0113), suggesting that IPSC charge transfer greatly increased by up/down-state induction and can suppress potentiated EPSCs in cortical neurons. However, neurons from het *Gabrg2^+/Q390X^* KI mice(het control) exhibited already large charge transfer ratio of eIPSC/eEPSC (4.11 ± 0.99, n=7 cells, n=3 mice) and up/down-state induction significantly decreased this ratio(Fig. 4B, 0.619 ± 0.259, n=10, n=3 mice, t-test p=0.0119), suggesting that IPSC charge transfer greatly reduced compared with potentiated EPSC charge transfer and creating an imbalance between eEPSCs and eIPSCs in cortical neurons from het *Gabrg2^+/Q390X^* KI mice. Similar as the charge transfer ratios, the ratios of eIPSC peak (holding at 0 mV) to eEPSC peak (holding at chloride reversal potential −89.1 mV) showed similar results (Fig. 4C/D)(wt control[n=7 cells] *vs* wt up/down[n=8 cells] from 1.02 ± 0.20 to 2.65 ± 0.49, n=2 mice each, t-test p = 0.012; het control[n=8 cells] *vs* het up/down[n=6 cells] from 4.38 ± 0.73 to 0.64 ± 0.227, n=2 mice each, t-test p = 0.0012), indicating that following up/down induction, eIPSCs in neurons from het *Gabrg2^+/Q390X^* KI mice were not able to balance the potentiated eEPSCs.

**Figure 4.**
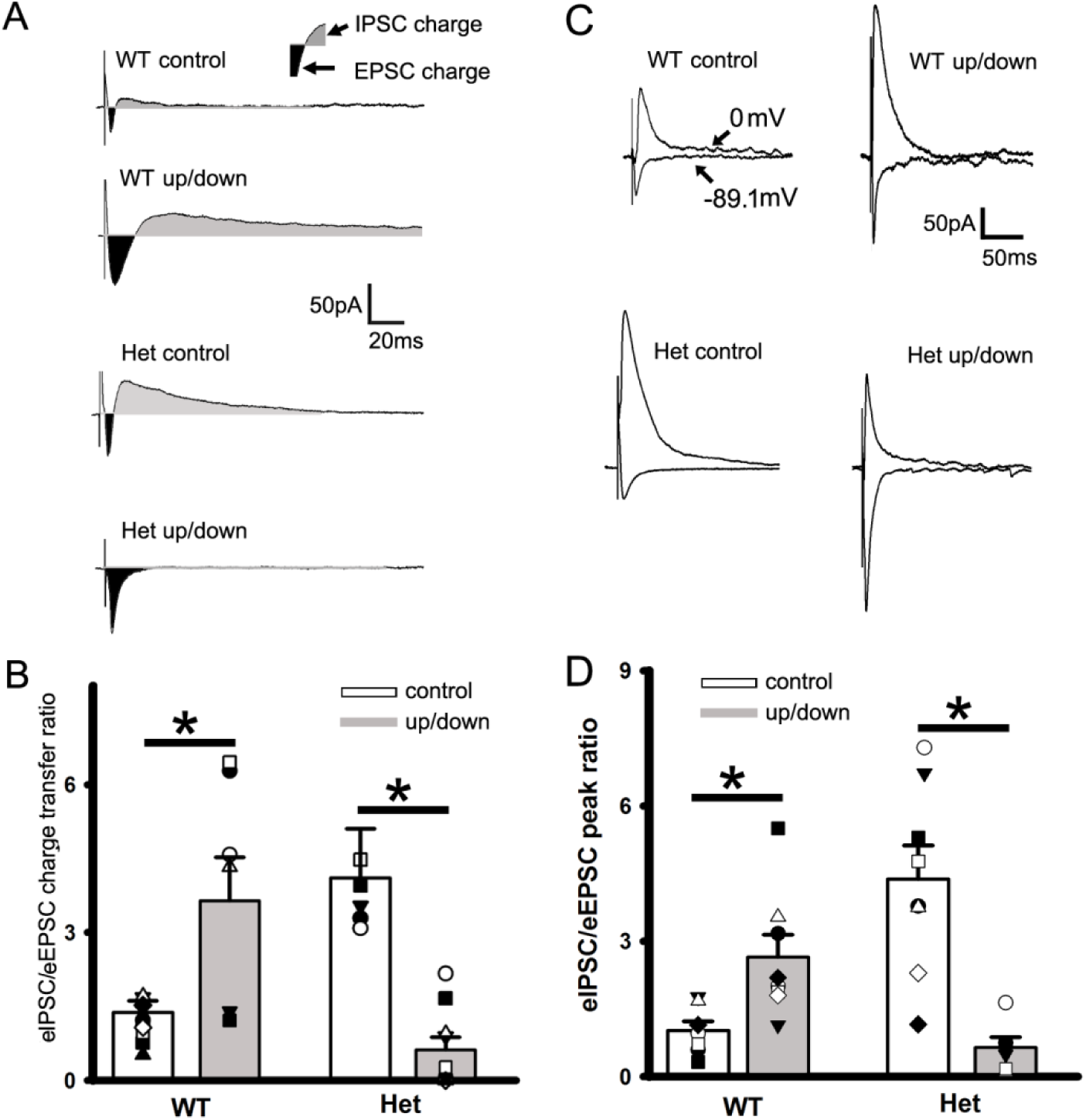
Balanced evoked(e) IPSCs/EPSCs following up/down-state induction in cortical neurons from wt littermates, while imbalanced eIPSCs/eEPSCs from het *Gabrg2^+/Q390X^* KI mice. A. Representative traces of paired/simultaneous eEPSCs and eIPSCs in neurons (clamped at −40mV) from wt control/wt up/down, het *Gabrg2^+/Q390X^* control/het *Gabrg2^+/Q390X^* up/down groups. The dark areas between inward current traces and baselines represent the excitatory current charges and the gray areas between outward current traces and baselines represent the inhibitory current charges. Stimulus artifacts are also shown right before eEPSCs. Scale-bars are indicated as labeled. B. Summarized data for these groups regarding eIPSC/eEPSC ratios (wt control n=11, n=3 mice and wt up/down n=7 cells, n=3 mice, t-test p=0.0113; het control n=7 cells, n=3 mice and het up/down n=10, 3 mice, t-test p=0.0119). Vertical Box and bars are averaged summary data (* mean significant difference). C. Representative traces of paired/simultaneous eEPSCs (holding at −89.1 mV) and eIPSCs (holding at 0 mV) in neurons from wt control/wt up/down, het *Gabrg2^+/Q390X^* control/het *Gabrg2^+/Q390X^* up/down groups. Stimulus artifacts are also shown right before eEPSCs and eIPSCs. Scale-bars are indicated as labeled. D. Summarized data for these groups regarding eIPSC/eEPSC peak ratios for wt control(n=7 cells) *vs* wt up/down (n=8 cells)(n=2 mice each, t-test p=0.012) and for het control(n=8 cells) *vs* het up/down (n=6 cells) n=2 mice each, t-test p=0.0012).

### Impaired potentiation of inhibitory synaptic currents effectively prompts cortical neurons to generate action potentials from het *Gabrg2^Q390X^* KI mice

Neuronal action potentials(APs) were evoked by local stimulation and stimulus intensity was adjusted to have some AP failures. In cortical neurons from wt littermates, SWO-induction(0.5 Hz) decreased the success rates of evoked APs compared to pre-SWO baseline(Fig. 5A and C, from 55.10 ± 0.097% to 20.05 ± 0.09%, n=8 cells, n=5 mice, paired t-test p=0.005), compatible with post-up/down potentiated eIPSC charge transfer(also larger peak ratio eIPSC/eEPSC) in neurons from wt littermates. In contrast, in cortical neurons from het *Gabrg2^+/Q390X^* KI mice, SWO induction significantly increased success rate of evoked APs compared to pre-SWO baseline(Fig. 5B-C, from 30.89 ± 10.16% to 65.10 ± 10.53%, n=8 cells, n=4 mice, paired t-test p=0.001), indicating that more neurons could fire action potentials following SWOs and contribute to synchronous neuron discharges *in vivo* in het *Gabrg2^+/Q390X^* KI mice, as SWOs *in vivo* are whole-brain macroscopic activity(Massimini, Huber, Ferrarelli, Hill, & Tononi, 2004; Volgushev, Chauvette, Mukovski, & Timofeev, 2006). This is compatible with post-up/down potentiated eEPSCs(not balanced by un-potentiated eIPSCs)(also smaller peak ratio eIPSC /eEPSC) in neurons from het *Gabrg2^+/Q390X^* KI mice. However, evoked AP number per each stimulus(during successful trials) did not reach any significant difference between wt littermates and het *Gabrg2^+/Q390X^* KI mice. In addition, there was no any significant difference in neuronal input-output curves regarding action potential generation between neurons from wt littermates and het *Gabrg2^+/Q390X^* KI mice(supplemental Fig. S4A-C, wt n=5 cells, n=2 mice and het n=6, n=3 mice, two-way ANOVA, p = 0.989).

**Figure 5.**
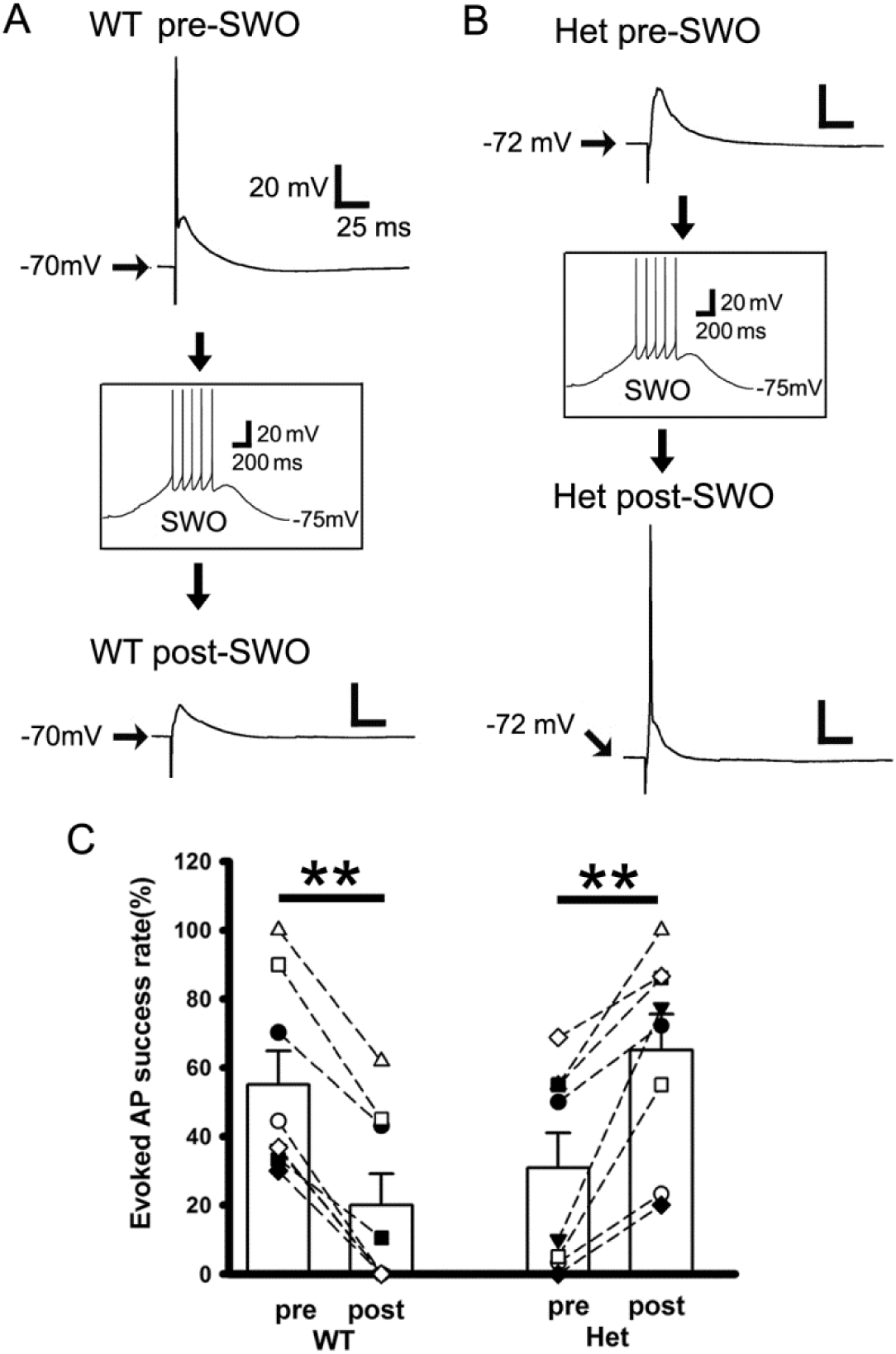
Impaired potentiation of inhibitory synaptic currents following SWOs more successfully prompts neurons to generate action potentials from het *Gabrg2^+/Q390X^* KI mice. Panels A and B are representative traces of pre-SWO (top) and post-SWO(lower) evoked APs or EPSPs in neurons from wt littermate mice(panel A) and het *Gabrg2^+/Q390X^* KI mice (panel B). Top panel(A or B) and lower(A or B) are two individual traces from the same neurons being recorded with successfully evoked APs or postsynaptic potentials. The middle insets(A or B) show one SWO induction from rest membrane potentials (−75mV). Scale bars are indicated as labeled. Panel C shows summary data for the success rate of evoked APs in wt littermate (n=8 cells, n=5 mice, paired t-test p=0.005) and het *Gabrg2^+/Q390X^* KI mice (n=8 cells, n=4 mice, paired t-test p=0.001). Each pair of pre- and post-SWO dots connected with a line corresponds to the same neuronal pre- and post-SWO AP success rates. Vertical Box and bars are averaged summary data (mean ± SEM for pre- and post-SWO evoked AP success rates).

### Induction of SWOs or up/down-states *in vivo* causes generalized epileptic SWDs in het *Gabrg2^Q390X^* KI mice

Using wt or het *Gabrg2^+/Q390X^* KI mice with Thy1-halorhodopsin expression and intracortical stimulation through planted tungsten electrodes within primary somatosensory cortex, we could induce SWOs or up/down-state alteration(0.5 Hz) *in vivo*. Electrical currents(300-400 pA used *in vivo*) were chosen to avoid kindling effect(kindling currents are within uA range, supplemental Fig. S5, n = 3 mice each)(Lothman, Bertram, Kapur, & Stringer, 1990; Racine, 1972a, 1972b) and an electrical duration of 20 ms was used to simulate sleep spindle duration during neuron up-state(Kandel & Buzsaki, 1997). Cortical neuron down-state were controlled by laser delivery (590 nm) to activate halorhodopsin. With enough intensity of laser delivered, neurons could be hyperpolarized by 5-10 mV from resting membrane potentials(supplemental Fig. S6 inset). After SWOs or up/down-state induction in neurons within *ex-vivo* brain slices(both wt and het *Gabrg2^+/Q390X^* KI mice), sEPSCs in cortical neurons increased their amplitudes compared to baseline(wt n=3 cells and het *Gabrg2^+/Q390X^* KI n=4 cells, total 5 mice, [wt and het sEPSC data together] sEPSC amplitude from 15.75 ± 1.95 to 22.30 ± 3.24 pA, paired t-test p=0.011), not sEPSC frequency(from 1.54 ± 0.71 to 1.30 ± 0.34 Hz, paired t-test p=0.725) and sEPSC decay τ(from 10.13 ± 1.92 to 44.75 ± 15.30 ms, paired t-test p=0.078), indicating that this laser delivery protocol could be used for SWO or up/down-state induction *in vivo*. Moreover, sIPSCs in cortical neurons within *ex vivo* brain slices from wt littermates, but not het *Gabrg2^+/Q390X^* KI mice, could be potentiated by laser-generated SWO or up/down induction(supplemental Fig. S6, wt sIPSC amplitude increased from 24.92 ± 3.43 to 33.47 ± 4.37 pA, n=5 cells, n=4 mice, paired t-test p=0.023; het sIPSCs were not significantly different from 15.92 ± 3.46 to 13.66 ± 2.74 pA, n=6 cells, n=4 mice, paired t-test p=0.100). However there was no any significant changes in sIPSC frequency and decay τ between baseline and laser-generated up/down-states(wt [n=5 cells, n=4 mice] sIPSC frequency from 3.99 ± 1.05 to 8.86 ± 2.76 Hz, paired t-test p=0.214; sIPSC decay τ from 15.17 ± 3.87 to 39.07 ± 11.04 ms, paired t-test p=0.085) (het [n=6 cells, n=4 mice] sIPSC frequency from 5.39 ± 2.00 to 2.64 ± 0.88 Hz, paired t-test p=0.210; sIPSC decay τ from 13.60 ± 7.84 to 15.92 ± 3.76 ms, paired t-test p=0.715).

In wt littermates, with laser delivery (intensity determined by those used in *ex vivo* brain slices) and intracortical stimulation to induce SWOs or up/down-states *in vivo*(0.5 Hz for 10min) (Fig. 6B), we observed only short EEG (band filtered between 0.1∼100 Hz) oscillation activity accompanied by a few multi-unit activity events (band filtered between 300∼3K Hz). Compared to pre-SWO baseline EEG/multi-unit activity in wt littermates, post-SWO EEG/multi-unit activity were similar and no significant changes were observed(Fig. 6 A and C-D). In contrast, during pre-SWO baseline recordings in het *Gabrg2^+/Q390X^* KI mice(Fig. 6E), EEG/multi-unit activity exhibited bilateral epileptic SWDs (accompanied by animal immobility) and slightly prolonged multi-unit burst activity events (Fig. 6E). During laser-generated SWOs or up/down-states *in vivo* in het *Gabrg2^+/Q390X^* KI mice(Fig. 6F), epileptic SWDs were observed and multi-unit activity became longer (Fig. 6F)(compared to non-SWD EEG and shorter multi-unit activity in wt littermates[Fig. 6B] during SWO induction *in vivo*). Furthermore, following laser-generated SWO or up/down-state induction *in vivo* in het *Gabrg2^+/Q390X^* KI mice(Fig. 6G), more epileptic SWDs(accompanied by animal immobility) and longer multi-unit activity events appeared (Fig. 6G-H), compared to pre-SWO baseline EEG/multi-unit activity(Fig. 6E). Strikingly, incidence of post-SWO epileptic SWDs in het *Gabrg2^+/Q390X^* KI mice exhibited a peak around 10-15 min along their time-course(Fig. 6H, gray for post-SWO SWDs), which proximately coincided the post-SWO sEPSC maximal enhancement(around 10∼15 min) in cortical neurons from het KI mice(Fig. 1E), suggesting that post-SWO unbalanced/potentiated sEPSCs (due to no potentiated sIPSCs) likely caused epileptic SWDs in het *Gabrg2^+/Q390X^* KI mice. Our summary SWD data indicated that in het *Gabrg2^+/Q390X^* KI mice, all the number(#) of post-SWO epileptic SWDs, total SWD duration and averaged single SWD duration significantly increased, compared to pre-SWO baseline SWDs(Fig. 6 I-K) (het *Gabrg2^+/Q390X^* KI mice [n=7 each], SWD # from 98.95 ± 14.29 to 201.01 ± 11.26/hour(hr), paired t-test p=0.001; total SWD duration from 317.71 ± 55.95 to 1331.56 ± 103.98 s/hr, paired t-test p=0.0008; averaged single SWD duration from 2.52 ± 0.24 to 6.26 ± 0.97 s, paired t-test p=0.005). In addition, in wt littermates, we did not observe significant difference in SWD #, total SWD duration and averaged single SWD duration compared to the basline(wt mice [n=6 each], SWD # from 8.5 ± 1.62 to 16.83 ± 3.45/hr, paired t-test p=0.104; total SWD duration from 19.74 ± 4.19 to 54.66 ± 12.3 s/hr, paired t-test p=0.088; averaged single SWD duration from 2.70 ± 0.46 to 3.34 ± 0.60 s, paired t-test p=0.165).

**Figure 6.**
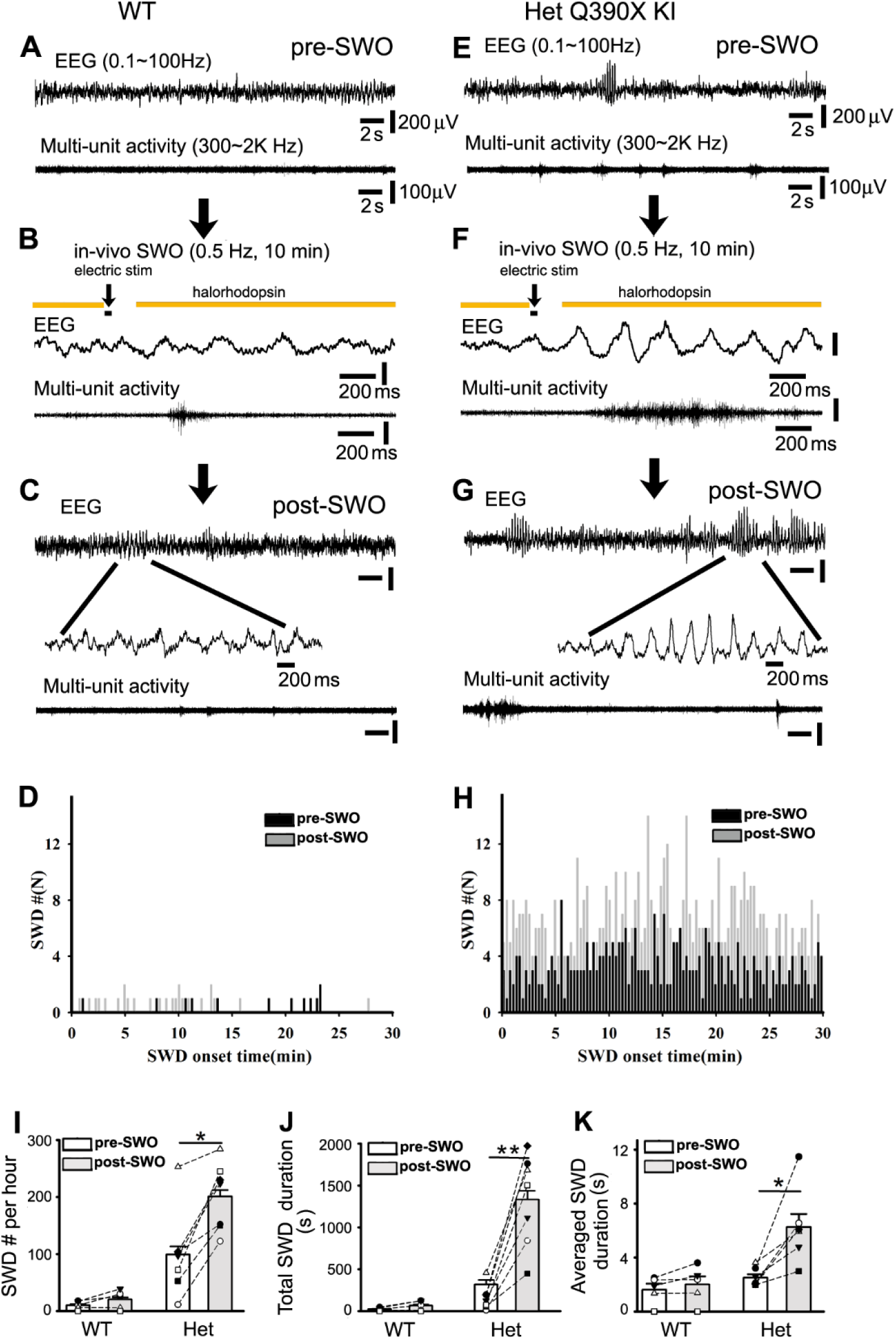
Induction of SWOs or up/down-states *in vivo* triggers epileptic SWDs generation in het *Gabrg2^+/Q390X^* KI mice. Panels A(wt) and E(het *Gabrg2^+/Q390X^* KI) are representative traces for pre-SWO simultaneous EEG(top) and multi-unit(below) recordings (30 s long). Panels B(wt) and F(het *Gabrg2^+/Q390X^*) show the time course of SWO or up/down-state induction *in vivo* with yellow color indicating laser delivery, and black bar for intracortical stimulation (20 ms). Below are representative traces(2 s long) for simultaneous EEG/multi-unit activity during SWO induction *in vivo*. Panels C(wt) and G(het *Gabrg2^+/Q390X^*) show representative traces for post-SWO simultaneous EEG(top) and multi-unit(below) recordings. Scale bars (panels A, C, E and G) are similarly indicated as labeled except time-scale bars in panels B and F. The downward arrows indicate experiment sequential steps. Panels D(wt) and H(het *Gabrg2^+/Q390X^*) show graphs of pre-(black) and post-SWO (gray) epileptic SWD onset time distribution. SWD events were obtained during 30 min right before and after SWO induction (1 hour(hr)). Panels I-K show summary data for SWD # per hr, total SWD duration and averaged single SWD duration for wt (blank bars) and het (gray bars) (het *Gabrg2^+/Q390X^* KI mice n=7 each, SWD #, paired t-test p=0.001; total SWD duration, paired t-test p=0.0008; averaged single SWD duration, paired t-test p=0.005) (wt mice n=6 each, SWD #, paired t-test p=0.104; total SWD duration, paired t-test p=0.088; averaged single SWD duration, paired t-test p=0.165).

Additionally, high-frequency synaptic activity were recorded with tungsten electrodes (band filtered 400∼800Hz)(Lasztoczi, Antal, Nyikos, Emri, & Kardos, 2004) and always preceded epileptic SWDs in het *Gabrg2^+/Q390X^* KI mice(supplemental Fig. S7)(wt pre-SWO 246.25 ± 31.12 ms [n=5 mice] *vs* het KI pre-SWO 380.38 ± 37.88 ms [n=6 mice], t-test p=0.026), implying that high-frequency synaptic activity could cause these epileptic SWDs. Following SWOs or up/down-state induction *in vivo*, the precedent high-frequency field potential activity became longer in het *Gabrg2^+/Q390X^* KI mice(supplemental Fig. S7C)(averaged episode duration from pre-SWO 380.38 ± 37.88 to post-SWO 491.16 ± 53.43 ms, paired t-test p=0.013, n=6 mice), while there was not significant changes in wt mice(from pre-SWO 246.25 ± 31.12 to post-SWO 224.42 ± 54.24 ms, paired t-test p=0.422, n=5 mice). This implies the involvement of SWO-induced potentiation of excitatory synaptic currents in neurons from het *Gabrg2^+/Q390X^* KI mice.

Due to very low incidence of generalized tonic-clonic seizures(GTCSs) and the limited experimental duration of pre-SWO (30 min) and post-SWO (1 hr), we could not firmly determine the effect of SWO induction on GTCSs. Also, induction of SWO *in vivo* did not cause any changes in juvenile myoclonic seizures, which is reasonable because juvenile myoclonic seizures are generated in frontal motor cortex(F. Arain et al., 2015; Ding, Satish, Zhou, & Gallagher, 2019; O’Muircheartaigh et al., 2012) and our optogenetic manipulation was administrated in the posterior primary somatosensory cortex.

## Discussion

In this study we found that sleep-like SWOs(0.5 Hz) or up/down-state could induce synaptic plasticity in cortical pyramidal neurons(layer V) from wt littermates, and that the state-dependent potentiation of excitatory or inhibitory synaptic currents remained intact in cortical neurons. In contrast, only excitatory synaptic currents, but not inhibitory, could be potentiated in neurons from het *Gabrg2^+/Q390X^* KI mice. Similar to homeostatic synaptic potentiation in culture neurons, SWO-induced state-dependent potentiation of excitatory synaptic currents in cortical neurons depended on low-level calcium, βCaMKII activation and retinoid acid synthesis, but not activation of L-type calcium channels. Moreover, similar to SWOs, physiologically similar up/down-state induction induced state-dependent potentiation of synaptic currents in cortical neurons. Following SWO induction, cortical neurons from het *Gabrg2^+/Q390X^* KI mice exhibited unbalanced evoked excitatory synaptic currents (smaller eIPSC/EPSC charge transfer ratios and smaller eIPSC/eEPSC peak ratios), prompting neurons to effectively generate action potentials, while neurons from wt littermates exhibited less action potential firings. With *in vivo* up/down-state induction by optogenetic manipulation(simulating *ex vivo* SWOs), epileptic SWDs could be elicited with significantly higher incidence and longer duration in het *Gabrg2^+/Q390X^* KI mice, not in wt littermates. These epileptic SWDs were accompanied by animal immobility and other epileptic behaviors, suggesting that SWOs or up/down-states *in vivo* can trigger epileptic seizure onset in IGE patients while they are in NREM sleep or quiet-wakefulness state.

Cortical neurons *in vivo* maintain stable activity during development and sleep-wake period by homeostatic synaptic plasticity mechanism(Levenstein, Watson, Rinzel, & Buzsaki, 2017; Turrigiano & Nelson, 2004; Watson, Levenstein, Greene, Gelinas, & Buzsaki, 2016). Homeostatic synaptic plasticity of neurons within cultures *in vitro* has been induced by using TTX/NBQX(low-level neuronal activity) or picrotoxin(high-level neuronal activity) (Chen et al., 2014; Kurotani et al., 2008; Turrigiano, 2008), indicating that manipulating neuron activity can regulate synaptic strength to feedback-influence neuronal action potential firings and maintain neuronal activity stability. In epileptic brain slices after hypoxic seizures(high-level neuronal activity), synaptic strength is homeostatically scaled down(Sun, Kosaras, Klein, & Jensen, 2013) to restrain neuron hyperexcitability. However, whether more physiology neuron activity during sleep states *in vivo*(whether NREM or REM sleep state) can engage homeostatic synaptic plasticity in cortical neurons or not remains questionable (Chauvette, Seigneur, & Timofeev, 2012; Gonzalez-Rueda, Pedrosa, Feord, Clopath, & Paulsen, 2018; Levenstein, Buzsaki, & Rinzel, 2019; Levenstein et al., 2017; Liu et al., 2010; Watson et al., 2016). In our study, 0.5 Hz SWOs resemble *in vivo* EEG delta wave oscillation(0.5 Hz) during NREM sleep and neuron membrane potential alteration during 0.5 Hz SWO induction (Fig. 1 and 2) mirrors neuron up/down-states (low-level activity) during NREM sleep or quiet wakefulness(Petersen et al., 2003). Moreover, using fast perfusion with one modified ACSF (3.5 or 5 mM K^+^ and 1 mM Mg^2+^ and Ca^2+^), physiological-like up/down-states in neurons can be induced within *ex vivo* brain slices from both wt littermate and het *Gabrg2^+/Q390X^* KI mice (Fig. 3, 10 s long up-states), very similar to findings from other groups(Neske, 2015; Neske et al., 2015; Rigas & Castro-Alamancos, 2007; Sanchez-Vives & McCormick, 2000). Thus, our results suggest that similar to low-level activity manipulation for homeostatic synaptic potentiation in cultures(using TTX/NBQX), SWOs(0.5 Hz) or up/down-states *in vivo* can engage homeostatic potentiation(scaling up) of synaptic currents in cortical neurons(both sEPSC/sIPSCs from wt litteramtes and only sEPSCs from het *Gabrg2^+/Q390X^* KI mice). Supporting this argument, our pharmacology experiments further indicate that this SWO-induced state-dependent synaptic potentiation(sEPSCs) requires low-level calcium and βCaMKII activation, and that nifedipine greatly enhances this synaptic potentiation(supplemental Fig. S2). Moreover, consistent with previous studies(Li et al., 2019), suppressing of retinoid acid synthesis by DEAB completely blocks this synaptic potentiation induced by SWOs or up/down-state. In addition, although we did not test the pharmacology of sIPSC potentiation by SWOs in neurons(from wt litteramtes), one similar study by Kurotani et al.(Kurotani et al., 2008)(using similar 0.5 Hz SWOs) does show calcium/nifedipine dependence of sIPSC potentiation in cortical neurons. Other than the synaptic scaling up by low-level activity(0.5 Hz SWOs or up/down-state), high-level activity with 50 Hz oscillation activity in cortical neurons can induce synaptic scaling-down of sEPSCs, which is very similar to neuron hyperexcitability state during hypoxic epilepsy(Sun et al., 2013). In a similar way, high-level activity of 5 Hz oscillation activity in neurons suppresses sIPSCs, indicating that homeostatic plasticity of sEPSCs and sIPSCs in neurons may have different set points of activity-dependent homeostatic plasticity. All these together suggest that SWO-induced state-dependent synaptic potentiation may entail some classic homeostatic plasticity mechanisms, although they may not be the same(Li et al., 2019; Thiagarajan et al., 2002; Turrigiano, 2008; Turrigiano & Nelson, 2004).

In addition, SWO induction for 10 min in most cortical neurons(80-90%) is able to engage homeostatic synaptic plasticity, which is compatible with previous reports that 5-10(Frank, Kennedy, Goold, Marek, & Davis, 2006), 30 min(Hou, Gilbert, & Man, 2011) or longer time(within hours) manipulation of neuron activity(Aoto et al., 2008; Ibata et al., 2008; Sutton et al., 2006) can induce synaptic homeostasis or hemostatic synaptic plasticity. We did observe that in some neurons(less than 20-10%), SWO induction for 5-6 min still can engage homeostatic synaptic plasticity, suggesting that engagement of homeostatic synaptic plasticity can be a fast process in our study. Moreover, the temporal period of SWO or up/down activity applied in this study are within temporal ranges of up/down state activity of cortical neurons *in vivo* and EEG SWO duration during slow-wave sleep, making our low-level activity manipulation for SWOs and up-down state more physiologically relevant *in vivo*.

Interestingly, induction of SWOs (0.5 Hz) or up/down-states is similarly effective to induce homeostatic potentiation of both excitatory and inhibitory currents in neurons (Fig. 1 and 2) as neuron activity manipulation(sinusoidal currents) *in vitro*(Chen et al., 2014; Kurotani et al., 2008; Turrigiano, 2008). This indicates that neuron excitatory and inhibitory currents are kept in a balance during different neuron activity states, as some studies have suggested *in vivo*(Adesnik & Scanziani, 2010; Dehghani et al., 2016; Moore, Weible, Balmer, Trussell, & Wehr, 2018; Shu et al., 2003; Takahashi, Kobayashi, Ishikawa, & Ikegaya, 2016). Our observations of eIPSC/eEPSC ratios in wt littermates are consistent with this dynamic balance mechanism(Fig. 4), similar to other findings(Dehghani et al., 2016; Haider et al., 2006). During normal physiology states (Fig. 4 het control), eEPSCs and eIPSCs(before up/down-induction) in neurons can be able to maintain a balance in het KI mice as wt littermates. However, during slow-wave sleep, up/down-state induction in neurons from het *Gabrg2^+/Q390X^* KI mice can potentiate sEPSCs, not sIPSCs (Fig. 2D-E), and at same time cause smaller eIPSC/eEPSC ratios(Fig. 4, het up-down), indicating that the balance between eEPSCs and eIPSCs has been state-dependently disrupted to cause seizures in IGE models(Bateup et al., 2013; Huguenard & McCormick, 2007; McCormick & Contreras, 2001). To the best of our knowledge, this is the first time that the impaired SWO-induced potentiation of inhibitory synaptic currents in cortical neurons from het *Gabrg2^+/Q390X^* KI mice can create one state-dependent imbalance between potentiated EPSCs and un-potentiated IPSCs, which likely prompts neuron synchronous firing(Fig. 5)[considering that SWOs *in vivo* is a whole-brain macroscopic activity condition(Massimini et al., 2004; Volgushev et al., 2006)]. In accordance with this, incidence of post-SWO epileptic SWD peak onset time(Fig. 6H het *Gabrg2^+/Q390X^*, gray trace, post-SWO) almost parallels the potentiated sEPSC amplitude peaks following SWOs(Fig. 1E het *Gabrg2^+/Q390X^*), indicating that SWO-induced imbalance between potentiated sEPSCs and un-potentiated sIPSCs eventually triggers epileptic SWDs and other epileptic motor behaviors(Fig. 6H het *Gabrg2^+/Q390X^* post-SWO). Moreover, post-SWO high-frequency field potential activity (400∼800 Hz, due to synaptic activity) becomes significantly longer to drive subsequent epileptic SWDs(supplemental Fig. S7). This is supported by one epileptic study *in vitro* that high-frequency synaptic activity causes epileptic activity(Lasztoczi et al., 2004). In addition, the impaired potentiation of sIPSCs in our het *Gabrg2^+/Q390X^* KI mice is very likely due to impaired GABAergic receptor forward trafficking(Kang et al., 2010; Zhou et al., 2013), which is very similar to the homeostatic mechanism proposed by Kurotani et al.(Kurotani et al., 2008).

IGE patients often have a genetic basis for their seizure activity generation(Allen et al., 2013; Striano & Zara, 2017). However, their seizure onset remains unpredictable. Recent review papers in one journal (review articles in Epileptic disorders, volume 21 supplement 1, 2019) discussed the slow-wave sleep association with seizure incidence. In addition, other acquired epileptic patients also show SWO activity preference before seizure onset(Lagarde et al., 2019). However, to our knowledge, there are no any mechanisms to explain this SWO preference in neuroscience and epilepsy fields. Our findings with SWO-induced seizure onset mechanisms (through state-dependent impairment of GABAergic synaptic currents in cortical neurons) offer one plausible/likely operational mechanism for epileptic SWD preferential incidence during NREM sleep and quiet-wakefulness state, which both generate SWOs in brain(Ahmed & Vijayan, 2014; F. Arain et al., 2015; Bagshaw et al., 2014; Halasz et al., 2002; Ng & Pavlova, 2013; Shouse et al., 2004). Due to SWOs *in vivo* being synchronized in the whole-brain(Contreras, Destexhe, Sejnowski, & Steriade, 1996; Massimini et al., 2004; Volgushev et al., 2006), SWO-induced unbalanced/potentiated EPSCs in cortical neurons of human IGE patients can likely synchronously occur in many epileptic foci within the whole brain and prompt simultaneous neuron firings to eventually trigger epileptic activity(Beenhakker & Huguenard, 2009). This may explain why IGE patients (including Dravet syndrome and absence epilepsy) are seizure-free during most active states and why seizure incidence frequency becomes much higher during NREM sleep and quiet-wakefulness state(Bagshaw et al., 2014; Halasz et al., 2013; Verbeek et al., 2015). And this may also apply to other acquired seizure onset during NREM preference patterns(Lagarde et al., 2019) if GABAergic synaptic potentiation is impaired.

In conclusion, sleep-like SWOs could induce state-dependent synaptic potentiation in cortical pyramidal neurons, which shares some homeostatic plasticity mechanisms, and potentiated excitatory and inhibitory synaptic currents remained in balance from wt littermates. However, only excitatory, but not inhibitory, synaptic currents can be potentiated in neurons from het *Gabrg2^+/Q390X^* KI mice, indicating that the dynamic balance between excitatory and inhibitory synaptic currents has been disrupted in het *Gabrg2^+/Q390X^* KI mice and neurons become more successful in generating action potentials. Using *in vivo* SWO or up/down-state induction by optogenetic laser delivery/intracortical stimulation, epileptic SWDs in het *Gabrg2^+/Q390X^* KI mice can be elicited with significantly higher incidence frequency and longer duration, accompanied by animal immobility and other epileptic behaviors. This suggests that SWOs *in vivo* may effectively trigger epileptic seizure onset in IGE patients during NREM sleep or quiet-wakefulness state, potentially leading to new translational therapeutic treatments of IGE.

## Materials and Methods

### Mouse brain slice preparation

All procedures were in accordance with guidelines set by the Institutional Animal Care and Use Committee of Vanderbilt University Medical Center. The brain slice preparation method used in this study has been described in previous studies(Zhou et al., 2013; Zhou, Lippman, Sun, & Jensen, 2012). Wt littermate and het *Gabrg2^+/Q390X^* KI mice aged P60-120 (both male and female) were used in this study. Transgenic animals expressing halorhodopsin were also used when necessary by crossing our het *Gabrg2^Q390X^* KI mice with Thy1-eNpHR2.0-EYFP transgenic mice(#012334, The Jackson Laboratory). Mice were deeply anesthetized and decapitated. Then coronal brain slices (300-320 µm thickness) containing somatosensory cortex were prepared with a vibratome(Leica VT 1200S, Leica Biosystems Inc) in a sucrose-based ice-cold solution (containing [in mM]): 214 sucrose, 2.5 KCl, 1.25 NaH_2_PO_4_, 0.5 CaCl_2_, 10 MgSO_4_, 24 NaHCO_3_, and 11 D-glucose, pH 7.4) and were incubated in a chamber at 35-36°C for 40 min with continuously oxygenated ACSF (see below for components). Finally slices remained at room temperature for at least 1 hour before electrophysiological recordings at 32°C.

### Brain slice electrophysiolog

Whole-cell patch-clamp recordings (voltage-or current-clamp) were made from somatosensory cortex layer V pyramidal neurons by using a Nikon infrared/DIC microscope(Eclipse FN1, Nikon Corp. Inc., Melville NY), and slices were continuously superfused (flow speed 1-1.5 ml/min) with ACSF (containing [in mM]: 126 NaCl, 2.5 KCl, 1.25 NaH_2_PO_4_, 2 CaCl_2_, 2 MgCl_2_, 26 NaHCO_3_, 10 D-glucose, pH 7.4) bubbled with 95% O_2_/5% CO_2_. Filled electrodes had resistances of 2∼5 MΩ with one internal solution for sEPSCs (consisted of [in mM]: 120 K-gluconate, 11 KCl, 1 MgCl_2_, 1 CaCl_2_, 0.6 EGTA, 10 HEPES, 2 Na-ATP, 0.6 Na-GTP, 10 K-creatine-phosphate, pH 7.3(Huguenard & Prince, 1994; Schofield, Kleiman-Weiner, Rudolph, & Huguenard, 2009)). sEPSCs were recorded at a holding potential −55.8mV (Cl^-^ reversal potential). The internal solution for sIPSCs contained (in mM): 65 K-gluconate, 65 KCl, 10 NaCl, 5 MgSO_4_, 0.6 EGTA, 10 HEPES, 2 Na-ATP, 0.6 Na-GTP, 10 K-creatine-phosphate, pH 7.3. sIPSC recorded at a holding potential −60 mV with AMPAR/kainate receptors being blocked (20 µM NBQX in ACSF)(Kurotani et al., 2008). For action potential(AP) recordings, the same internal solution for sEPSC recordings was used. A concentric bipolar tungsten electrode for stimulation was placed into the neighboring area around neurons being recorded. The stimulus pulses were 0.1 ms duration and stimulus intensity was adjusted to have some AP evoked failures, and the same stimulus intensity was repeated for 30-40 times with a 20-30 s interval during baseline and post-SWO period. Slow-wave oscillations (SWOs, 0.5Hz) were induced by injecting sinusoidal currents (200-300 pA at 0.5 Hz) into cells(current-clamp mode) to cause neuron membrane potential oscillating between membrane potential (around −75 mV) and −50 mV (with 4-5 AP firings) for 10 min. For some experiments, SWOs were also induced by injecting currents (150-300 pA, 200-300 ms) into cells (in current-clamp) along with 589 nm laser delivery to activate halorhodopsin for 10 min and these data with laser delivery in *ex vivo* slices were combined with experiments using SWO induction. For some experiments, neuron up/down-states were induced by using one modified ACSF (containing (in mM) 3.5 or 5 KCl, 1 Ca^2+^, 1 Mg^2+^ and 3.5 µM carbachol, other components were same as regular ACSF components) and fast flow perfusion (6-7 ml/min, at 32∼33°C)(Hajos & Mody, 2009; Neske et al., 2015; Reid, Edmonds, Schurr, Tseng, & West, 1988; Sanchez-Vives & McCormick, 2000). Access resistance(Ra) was continuously monitored and recordings with Ra larger than 25 MΩ or 20% change were discarded. Neuron input-output curves of AP firings were measured by injecting step-currents into neurons with all synaptic currents blocked(20 μM NBQX, 100 μM D-AP5 and 60 μM picrotoxin in ACSF). For 4-(diethylamino)-benzaldehyde(DEAB, blocking retinoid acid synthesis) experiments, all incubation solution and superfusing ACSF contained 40 μM DEAB, and brain slices were maintained in these solutions for at least 1 hour before recordings. KN-93 (2 μM) (Hello Bio Inc., Princeton NJ) was dissolved in DMSO and added into pipette internal solutions. BAPTA-AM (15 μM) was administrated in brain slice incubation solution for at least 40 min before recordings. Nifedipine (20 μM) was dissolved in ACSF during experiments. All chemicals were purchased from Sigma-Aldrich Inc. except specifically stated.

Paired evoked(e) EPSCs and eIPSCs in neurons were simultaneously recorded with one Cs-based internal solution(containing [in mM]: 145 Cs-gluconate, 2 MgCl_2_, 0.5 EGTA, 10 HEPES, 2 Tris-ATP, 0.2 Na-GTP, 5 QX-314, pH 7.3(Nanou et al., 2018))(Cl^-^ reversal potential is −89.1 mV) to measure charge transfer of eEPSCs/eIPSCs with D-AP5(100 μM) in ACSF. All evoked synaptic currents were in linear ranges of stimulating input-output curves(same intensity for paired eEPSCs/eIPSCs) and showed smooth rising phase without overlapped multi-synaptic events. Recordings with same stimulus intensity(0.1 ms duration) were repeated 10 times (interval 20-30 s) to average to remove synaptic fluctuation. Instead using SWOs induction which could not induced by using Cs-based internal solutions, WT up/down and Het up/down-states were used with one modified ACSF (containing [in mM] 3.5 or 5 KCl, 1 Ca^2+^, 1 Mg^2+^ and µM carbachol) and fast flow perfusion (6-7ml/min, at 32-33°C). After up/down-state induction in brain slices was confirmed by recordings(current-clamp mode) with the above K-gluconate based internal solution for sEPSC plasticity, subsequently paired eEPSCs/eIPSCs in neurons(by using Cs-based solutions) were recorded at −40 mV(for both eEPSC/eIPSC charge transfer) or at −89.1 mV(for eEPSC peak) and 0 mV(for eIPSC peak). The ratios of inhibition and excitation were calculated as the ratios of area under curves for eEPSCs/eIPSCs or as the ratios of eIPSC to eEPSC peaks.

### Data collection and analysis

Data were collected using one multiClamp 700B amplifier and Clampex 10 software (Molecular Devices Inc., Union City, CA) and filtered at 2 kHz, and digitized at 20 kHz using a Digidata 1440A(Molecular Devices Inc., Union City, CA). Both sEPSCs and sIPSCs were analyzed with a threshold detection method (5-6 pA, 2.5X baseline RMS) using Clampfit 10.0 software program(Rakhade et al., 2012; Schofield et al., 2009; Zhou et al., 2012). pre- and post-SWO sEPSC or sIPSC consecutive sweeps (each sweep 5s long) from the same neurons were concatenated as one file to be analyzed by using same threshold setting for all synaptic events. All detected sEPSC and sIPSC events were checked by eye to ensure that their waveforms had normal rising and decaying phases. Then sEPSC and sIPSC histogram and cumulative distribution graphs were constructed with the Clampfit 10.0 software and Kolmogorov–Smirnov(K-S) non-parametric test was performed. All figures were prepared with Microsoft Excel, SigmaPlot/Stat and Adobe Photoshop softwares. Data were expressed as mean ± SEM(standard error of mean). Two-way ANOVA and Holm-Sidak test was used to compare wt and het neurons input-output of AP data.

### Mouse surgery and EEG/multi-unit recordings *in vivo*

With all procedures in accordance with guidelines set by the Institutional Animal Care and Use Committee of Vanderbilt University Medical Center, both wt littermate and het *Gabrg2^+/Q390X^* KI mice (expressing halorhodopsin, by crossing mice #012334 from The Jachson Laboratory(Gradinaru, Thompson, & Deisseroth, 2008; Zhao et al., 2008)) underwent brain surgery (anesthesia 1–3% isoflurane (vol/vol)) to plant three EEG screw electrodes (each for one hemisphere and one for grounding in cerebellum, #8201 Pinnacle Technology, Lawrence KS), one concentric bipolar tungsten electrode in somatosensory cortex (S1 cortex, depth 0.8∼1 mm in laminar V) and one fiber optic cannula (0.2-0.4 mm diameter, Thorlabs Inc., Newton, New Jersey) for laser light delivery *in vivo* (brain atlas coordinates: within somatosensory cortex range, anterior-posterior between −1.82 and −0.46 mm, midline-lateral between +2.0 and +4.0 mm reference to bregma, dorsal-ventral 0.7-1.1 mm reference to pia surface). The tungsten electrode tip was placed a slightly deeper than the optic cannula depth in S1 cortex to ensure that all neurons recorded were stimulated by laser delivered(589∼680nm). We also used one pair of unipolar tungsten electrodes with one (together with one fiber optic cannula) within S1 cortex and another in posterior brain (anterior-posterior −2.5-2.6 mm and midline-lateral 0-0.3 mm, no optic cannula at this location), with similar effects to that of concentric bipolar tungsten electrodes planted in somatosensory cortex. One EMG lead was inserted into the trapezius muscles to monitor mouse motor activity. After surgery, mice were continuously monitored for recovery from anesthesia and remained in the animal care facility for at least 1 week (normal wake/sleep circadian rhythm) before simultaneous EEG/EMG/multi-unit activity recordings *in vivo* during daytime(mouse sleep period). Tungsten electrode/optic cannula placement within S1 cortex were checked in mouse brains after euthanasia. Simultaneous EEG (two channels, band filtered at 0.1-100 Hz) and multi-unit recordings (one-channel, band filtered 300-2000 Hz) (both in current-clamp mode) data along with one channel EMG recordings(400 Hz) were collected by using two multiClamp 700B amplifiers (total 4 channels, Molecular devices Inc., Union City, CA) and Clampex 10 software (Molecular Devices Inc., Union City, CA), and digitized at 20 kHz using a Digidata 1440A.

The laser was delivered through an fiberoptic cable connected to the optic cannula, controlled by a DPSS laser (MGL-III-589-50 (50mW, Ultralazers Co., Inc) and the timing control of laser delivery was controlled by Clampex 10 software. Intracortical stimulations (300-400 pA, 20 ms) were applied through planted tungsten electrodes by Clampex 10 software (current-clamp mode). SWOs or up/down-states *in vivo* (0.5 Hz, for 10 min) were induced by alternating laser delivery(for hyperpolarizing down-state, 1800 ms) and no laser delivery[for up-state 200 ms, at the beginning of this up-state, electrical stimulations(simulating sleep spindle duration(Kandel & Buzsaki, 1997; Levenstein et al., 2017)) were applied]. The injected current amplitudes(pA) and duration were chosen to avoid kindling effect which needs larger currents within uA amplitude range(Lothman et al., 1990; Racine, 1972a, 1972b). Mouse epileptic behaviors were video-recorded, and synchronized with EEG recordings and Racine-scaled. Bilateral synchronous spike-wave discharges (SWD, 6-12 Hz) and slow SWD [SSWD, 3-6 Hz(Cortez, McKerlie, & Snead, 2001; Velazquez, Huo, Dominguez, Leshchenko, & Snead, 2007)] were defined as trains (>1 s) of rhythmic biphasic spikes, with a voltage amplitude at least twofold higher than baseline amplitude(F. M. Arain et al., 2012; Velazquez et al., 2007). All atypical and typical absence seizures and general tonic-clonic seizures(GTCS) started with SSWDs or SWDs, accompanied by characteristic motor behaviors consisting of immobility, facial myoclonus and vibrissal twitching. The SWD/SSWDs and animal behaviors were also checked and confirmed by persons blind to animal genotypes. The onset times of SWD or SSWDs were determined by their leading edge points crossing (either upward or downward) the precedent EEG baseline. High-frequency activity was obtained by post-experiment band-filtering(400-800Hz) tungsten activity originally recorded at 300-3K Hz *in vivo*. One threshold detection method (at least 2X baseline amplitude) was used to detect high-frequency activity events using Clampfit 10 software(Molecular Devices Inc., Molecular Devices Inc., Union City, CA) and their duration was analyzed. Any high-frequency activity associated with mobile motor behaviors (video monitored) was removed from this analysis.

## ACKNOWLEDGMENTS

Chengwen Zhou and Chun-Qing Zhang designed experiments; Chengwen Zhou performed experiments; Li Ding and Caitlyn M. Hanna provided animal care and surgery. Chun-Qing Zhang, Mackenzie Catron and Chengwen Zhou analyzed data; and Chengwen Zhou wrote the manuscript and discussed with Martin J. Gallagher and Robert L. Macdonald.

This work was supported by National Institutes of Health Grants NINDS R21NS096483-01 (Gallagher, Macdonald, Zhou[contact]) and R01NS107424-01(Zhou). We also thank National Natural Science Foundation of China to support Chun-Qing Zhang’s visiting scholarship (2013-2015 at Vanderbilt University Medical Center Dept. of Neurology) from Department of Neurosurgery, Xinqiao Hospital, Army Military Medical University, Chongqing, China.

