## supplemental figure and legends for "Impaired state-dependent potentiation of GABAergic synaptic currents triggers seizures in an idiopathic generalized epilepsy model"

**Supplemental Material**

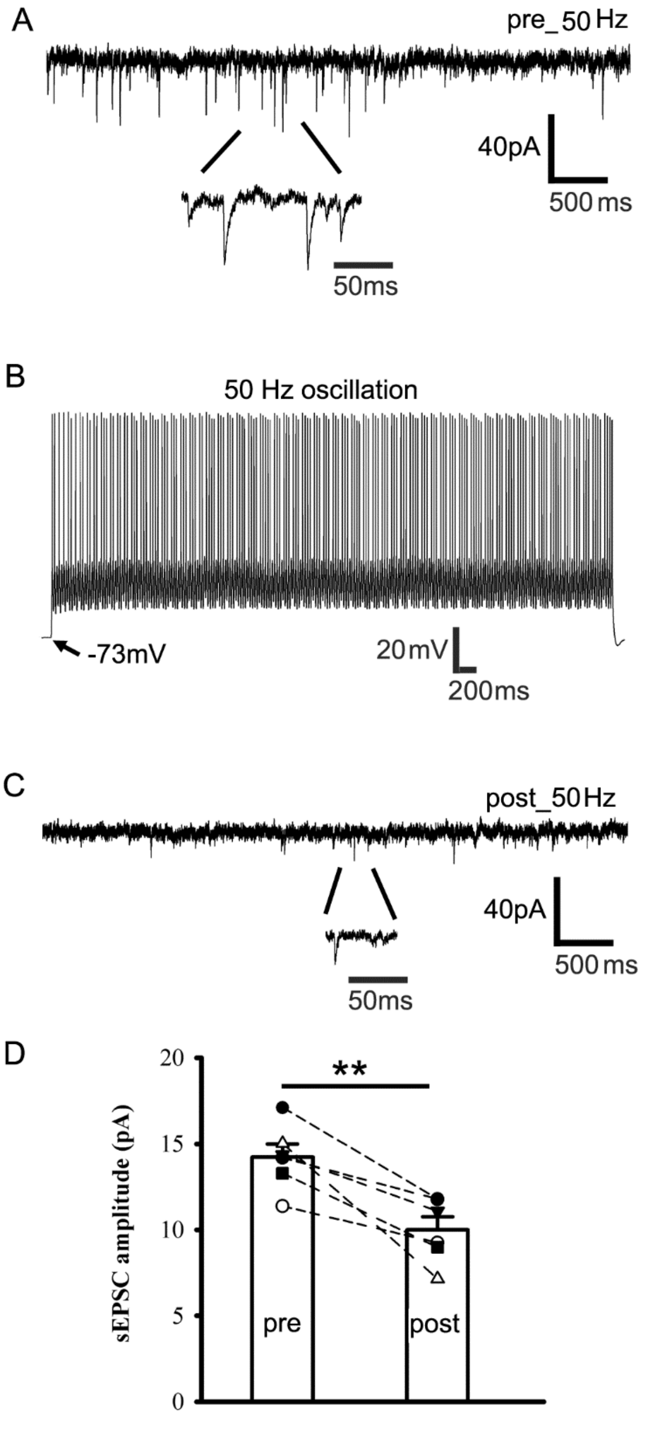


**Supplemental Figure S1. Elevated neuron activity (50 Hz) attenuates sEPSCs in cortical neurons from wt littermate or het *Gabrg2*$^{+/Q390X}$ KI mice.** Panels A and C show representative traces for pre- and post-50 Hz sEPSCs in cortical neurons.  Individual sEPSC events are expanded to show their rising and decaying phases. Panel B shows that 50 Hz oscillating activity was induced in cortical neurons for 10 min from resting membrane potential -72 mV. Scale bars are indicated as labeled. Panel D shows summary data for pre- and post-50 Hz sEPSCs (sEPSC amplitude n=6 cells, n=4 mice, paired t-test p=0.004).

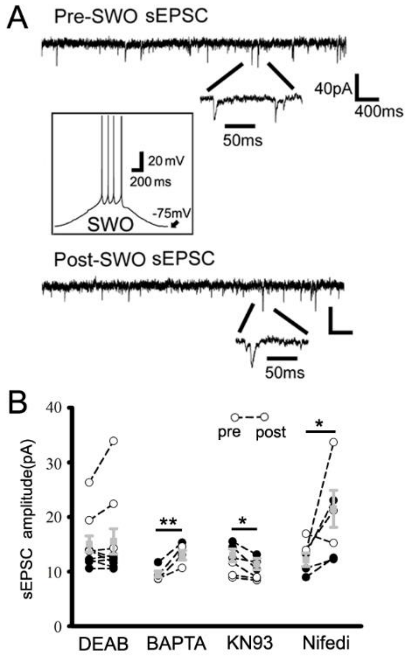


**Supplemental Figure S2. Pharmacology of SWO-induced potentiation of sEPSCs in cortical neurons.** Panel A shows representative traces for pre-(top) and post-SWO sEPSCs (bottom, for DEAB treatment) in cortical neurons.  Individual sEPSC events are expanded to show their rising and decaying phases. The inset shows SWO induction trace from resting membrane potential -75 mV. Scale bars are indicated as labeled. DEAB concentration used in ACSF was 40 µM. Panel B shows summary data for pre-SWO and post-SWO sEPSCs with DEAB treatment (n=10 cells, n=4 mice, paired t-test p=0.632), BAPAT-AM(15 µM, n=5 cells, n=3 mice, paired t-test p=0.006), KN93(2 μM, n=7 cells, n=3 mice, paired t-test p=0.013) and nifedipine(20 μM, n=7 cells, n=4 mice, paired t-test p=0.044).

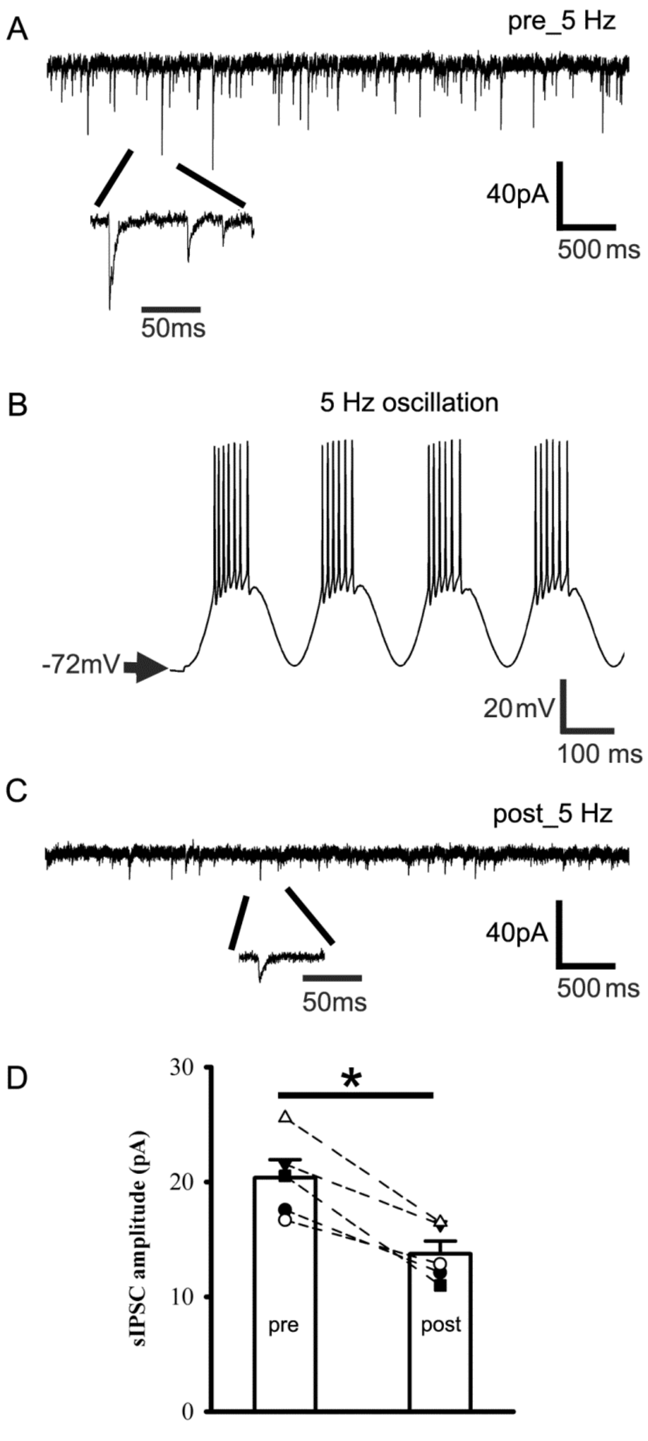

**Supplemental Figure S3. Elevated neuron activity (5 Hz) attenuates sIPSCs in cortical neurons from wt littermates.** Panels A and C show representative traces for pre- and post-5 Hz sIPSCs in cortical neurons. Individual sIPSC events are expanded to show their rising and decaying phases. Panel B shows that 5 Hz oscillating activity was induced in cortical neurons for 5-10 min from resting membrane potential -72mV. Scale bars are indicated as labeled. Panel D shows summary data for pre- and post-5 Hz sIPSCs (sIPSC amplitude n=5 cells, n=4 mice, paired t-test p=0.004 and sIPSC frequency n=5 cells, n=4 mice, paired t-test p=0.048, not shown in graph).

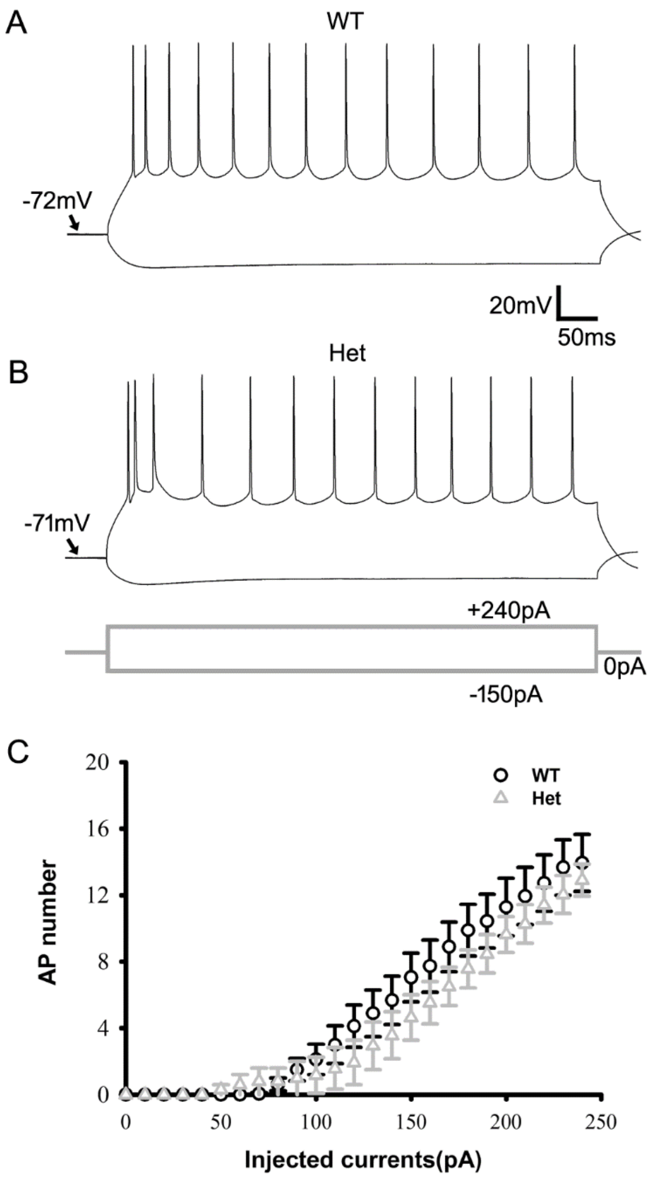


**Supplemental Figure S4. Cortical neuronal input-output curves of action potentials from wt littermate and het *Gabrg2*$^{+/Q390X}$ KI mice.** Panels A and B show representative traces action potential firing in layer V pyramidal neurons with different step-current injection(-150 to 240 pA) from wt littermate and het *Gabrg2*$^{+/Q390X}$ KI mice. Panel B (lower panel) shows the time course of injection step-currents. Scale bars are indicated as labeled. Panel C shows summary data (wt n=5 cells, n=2 mice and het *Gabrg2*$^{+/Q390X}$ KI n=6 cells, n=3 mice, two-way ANOVA p=0.989).

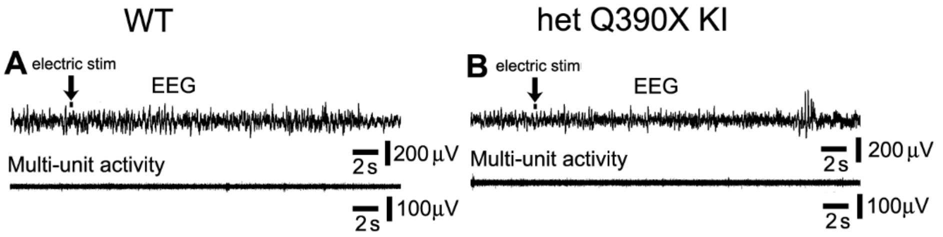


**Supplemental Figure S5. Intracortical stimulation alone does not cause any changes in cortical EEG/multi-unit activity in wt or het *Gabrg2*$^{+/Q390X}$ KI mice.** Panels A(wt) and B(het *Gabrg2*$^{+/Q390X}$ KI) are representative traces for simultaneous EEG (top) and multi-unit (below) recordings (30 s long) (by comparing to pre-SWO baseline in Fig. 6A/E). Intracortical stimulation (300-400 pA, 20 ms) was administrated through tungsten electrodes within somatosensory cortex (n=3 mice each). Scale bars are indicated as labeled.

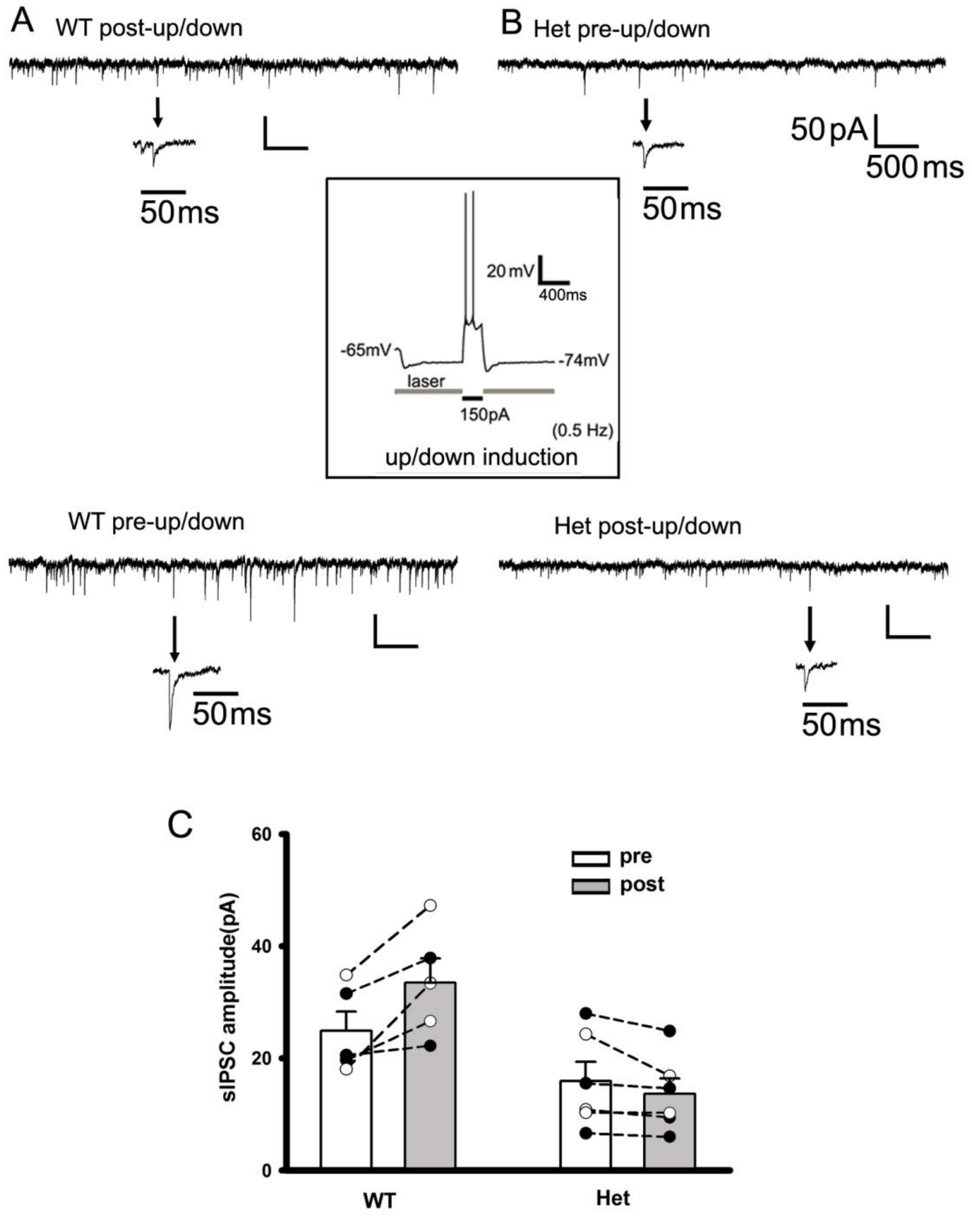


**Supplemental Figure S6. Up/down-state induction by laser delivery and current injection can potentiate sIPSCs in cortical neurons from wt littermates, not het *Gabrg2*$^{+/Q390X}$ KI mice.** Panel A shows representative traces of baseline sIPSCs(top) and post-up/down sIPSCs(lower) in wt cortical neurons. Panel B shows representative traces of baseline sIPSCs(top) and post-up/down sIPSCs(lower) in het *Gabrg2*$^{+/Q390X}$ cortical neurons. The middle inset shows the representative trace of up-down state induction with laser delivery(dropping from

-65 to -74 mV, down-state) and depolarization (around -52 mV, up-state, current injection 150 pA, 200 ms)(0.5 Hz for 10 min, similar as SWO induction). Individual sEPSC events are expanded to show rising and decaying phases. Scale bars are indicated as labeled. Panel C shows summary data for wt and het *Gabrg2*$^{+/Q390X}$ mice (wt n = 5 cells, n = 4 mice, paired t-test p = 0.023; n = 6 cells, n = 4 mice, paired t-test p = 0.100).

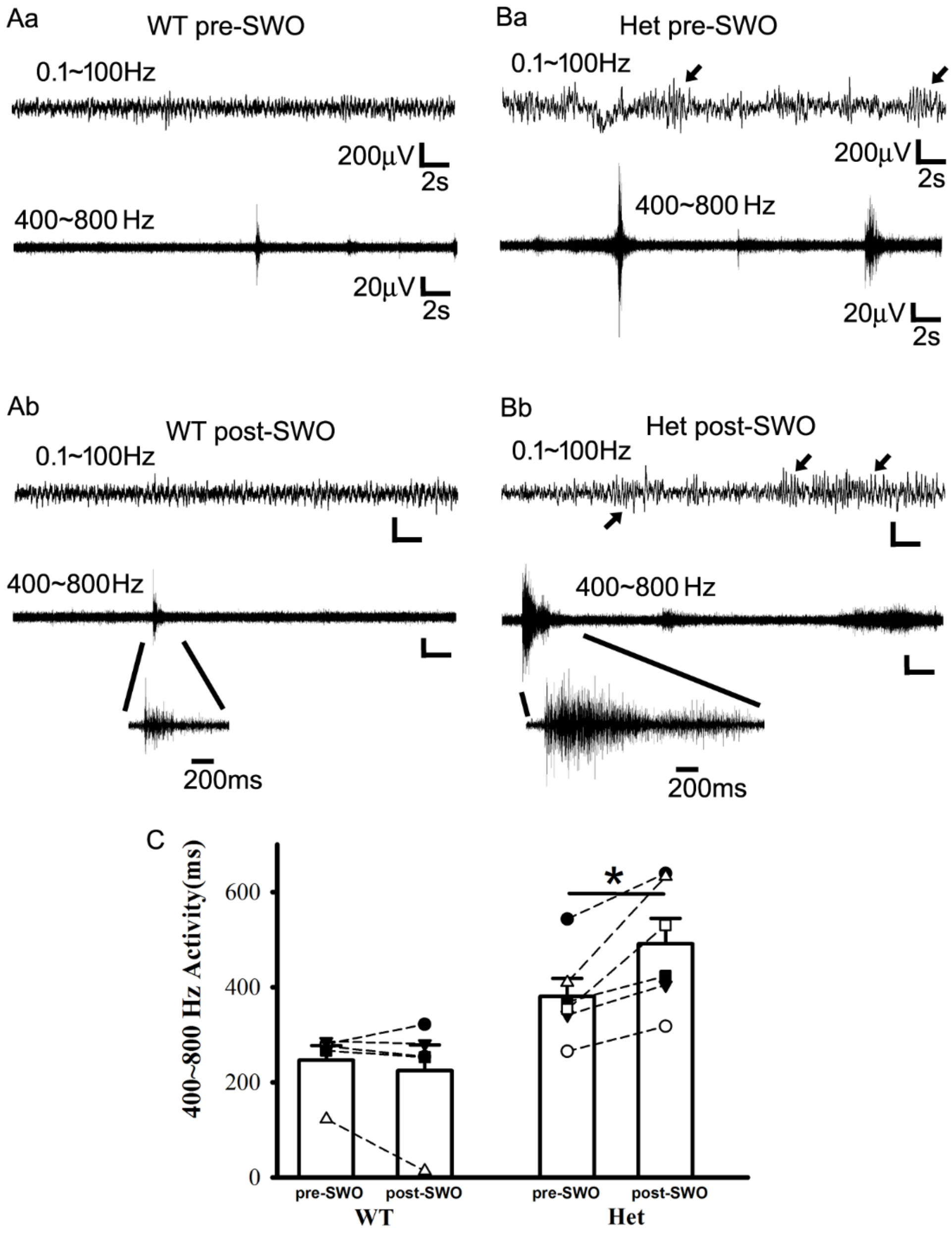

**Figure S7. High-frequency field potential activity precedes epileptic SWDs and becomes longer in het *Gabrg2*$^{+/Q390X}$ KI mice following SWO induction.** Panels Aa-b(wt) and Ba-b(het *Gabrg2*$^{+/Q390X}$) show representative traces for pre- and post-SWO tungsten activity *in vivo* (band-filtered between 400~800 Hz), along with simultaneous EEG activity (wt) or epileptic SWDs(het *Gabrg2*$^{+/Q390X}$). Scale bars are indicated as labeled. Arrows indicated epileptic SWDs in EEG recordings from het mice. Panel C shows summary data for pre- and post-SWO averaged duration of high-frequency activity from wt and het *Gabrg2*$^{+/Q390X}$ KI mice (wt n=5 mice, pre- and post-SWO paired t-test p=0.422 and het n=6 mice, pre- and post-SWO paired t-test p=0.013). The same symbols are used for data points of pre- and post-SWO high-frequency activity duration from the same mice.
